# LPS-induced delirium-like behavior and microglial activation in mice correlate with bispectral electroencephalography (BSEEG)

**DOI:** 10.1101/2024.09.27.615556

**Authors:** Tsuyoshi Nishiguchi, Kyosuke Yamanishi, Nipun Gorantla, Akiyoshi Shimura, Tomoteru Seki, Takaya Ishii, Bun Aoyama, Johnny R. Malicoat, Nathan James Phuong, Nicole Jade Dye, Takehiko Yamanashi, Masaaki Iwata, Gen Shinozaki

## Abstract

Delirium is a multifactorial medical condition characterized by impairment across various mental functions and is one of the greatest risk factors for prolonged hospitalization, morbidity, and mortality. Research focused on delirium has proven to be challenging due to a lack of objective measures for diagnosing patients, and few laboratory models have been validated. Our recent studies report the efficacy of bispectral electroencephalography (BSEEG) in diagnosing delirium in patients and predicting patient outcomes. We applied BSEEG to validate a lipopolysaccharide (LPS)-induced mouse model of delirium. Moreover, we investigated the relationship between BSEEG score, delirium-like behaviors, and microglia activation in hippocampal dentate gyrus and cortex regions in young and aged mice. There was a significant correlation between BSEEG score and impairment of attention in young mice. Additionally, there was a significant correlation between BSEEG score and microglial activation in hippocampal dentate gyrus and cortex regions in young and aged mice. We have successfully validated the BSEEG method by showing its associations with a level of behavioral change and microglial activation in an LPS-induced mouse model of delirium. In addition, the BSEEG method was able to sensitively capture an LPS-induced delirium-like condition that behavioral tests could not capture because of a hypoactive state.

## Introduction

Delirium is a dangerous disturbance in cognitive function and is especially prevalent among aged populations and those who have undergone major surgery^1^. Systematic reviews have shown delirium to be prevalent among 23% of generally hospitalized older adults, 20% of high-risk patients who have undergone major surgery, 25% of stroke patients, 59-88% of palliative care inpatients in the weeks prior to death, and 50-70% of mechanically ventilated patients^1^. Delirium has been associated with adverse patient outcomes, including increased rates of mortality, extended length of hospital stay, and lasting cognitive impairment^2–5^. Despite its pervasiveness and devastating consequences to patient health and quality of life, however, there have been few successful investigations identifying the underlying mechanisms of, and potential treatments for, delirium.

Currently, diagnosis primarily relies on clinical observation and expert opinion because there are few biological-quantitative monitoring methods^6–8^. For this reason, our group has worked on the quantitative identification of delirium in human patients using a novel bispectral electroencephalogram (BSEEG) method, which has been validated as an effective delirium detection tool in over 1000 patients^9–14^. While our BSEEG method is a promising clinical diagnostic tool, it is important to assess its utility in preclinical studies to lay the groundwork for a better understanding of the underlying molecular pathophysiology of delirium, a vital step toward the development of potential treatments.

In preclinical studies of delirium, a delirium-like state is usually induced by mock surgery or via injection of lipopolysaccharide (LPS), a bacterial toxin known to induce systemic- and neuro-inflammation characteristic of delirium^15,16^. Following the induction of a delirium-like state in model animals, investigators often rely on behavioral measures to characterize phenotypes associated with delirium^17,18^. While behavioral measures are crucial for gaining insight into the cognitive impairments associated with delirium, relying solely on qualitative behavioral outcomes often makes results hard to quantify and insufficient to precisely gauge the severity of the delirium pathophysiology in model animals.

In a previously published study, we created a novel BSEEG algorithm for the mouse model of delirium based on one designed to capture slow waves in humans^19^. We reported that intraperitoneal (i.p.) injection of LPS in mice induced a BSEEG score increase, capturing several delirium-like characteristics^19^. First, BSEEG scores in mice increased following LPS injection, reproducing the heightened BSEEG scores seen in patients with delirium following infection. Second, higher LPS injection doses correlated with higher BSEEG scores in a dose-dependent manner. This again mirrors the results of human studies demonstrating that the incidence and severity of delirium increased with severe infection^20^. Third, aged mice showed more pronounced increases in BSEEG scores than young mice following the same dose of LPS, reflecting the fact that elderly patients are more vulnerable to infection than younger patients^21^. Fourth, LPS administration disrupted the normal diurnal change in EEG, just as human patients with delirium lose their sense of day and night.

Thus, our BSEEG animal model of delirium shows promise as a reliable, quantifiable, and reproducible method for studying delirium in a preclinical setting. However, to validate the BSEEG method further, it is important to show that the BSEEG scores are representative of the cognitive impairments that characterize delirium, such as reduced attention, decreased consciousness, and loss of organized thinking. In this study, we investigate the relationship between BSEEG scores following LPS injections and a battery of behavioral tests in mice that have been used to characterize delirium-like behavior in literature^17,18^. We also aimed to test the relationship between the BSEEG score change in response to LPS exposure and microglial activation. One systematic review previously reported that there were various animal experiments related to the effects of systemic inflammation on the microglial and inflammatory response in the brain by LPS administration and that peripheral inflammatory stimulation may activate microglial cells in the brain, implicating an important role for microglia in sepsis-associated delirium^22^. One clinical case-control study also documented an inflammatory mechanism involved in delirium pathophysiology^23^. We hypothesize that an increase in the BSEEG score after LPS administration will be correlated with an increase in delirium-like phenotypes as measured by behavioral assays as well as microglial activation.

## Methods

### Animals and housing

All male C57BL/6J mice (young: 2-3 months and aged: 18-19 months) were purchased from Jackson Laboratory and were housed at the animal housing facility at Stanford University. Our study examined male mice because including females complicates age comparisons due to changing female hormone levels with age. Before the EEG head mount implantation surgery, 4-5 mice shared a cage with free access to food and water. After the surgery, each mouse was housed individually. All mice were randomly allocated to the saline and LPS groups. There were no significant differences in body weights between experimental groups (eFigure 1). The facility treated the mice to a 12:12 h light-dark cycle (7 am on/7 pm off) at a temperature of 20-26L and humidity of 30-60%.

### Study approval

The animal experiments were conducted following a protocol approved by an Institutional Animal Care and Use Committee (IACUC). At Stanford, the IACUC is known as Stanford’s Administrative Panel on Laboratory Animal Care (APLAC). The APLAC is accredited by the Association for Assessment and Accreditation of Laboratory Animal Care (AAALAC).

### Reagents and resources

All reagents and resources information are in eTable 1.

### Experimental procedure

#### Schedule

The full experimental schedule of EEG head mount implantation surgery, EEG recording, LPS injection, and behavioral tests is shown in eFigure 2.

##### Experiment 1

On day -14, C57BL/6J young mice (2-3 months old) underwent an EEG head mount implantation. The postoperative recovery period was 14 days. On day 1, we started EEG recording to measure the baseline BSEEG score for three days (days 1-3). On day 4, we conducted a battery of three types of behavioral tests (Buried Food Test (BFT), Open Field Test (OFT), and Y-maze test) that have been used with mouse models of delirium. These tests were conducted to establish baseline behavior 24 hours before LPS administration. On day 5, we administered saline or LPS (0.5 mg/kg and 1.0 mg/kg) to young mice by i.p. injection, and behavioral tests were repeated 6 hours after injection (n□=□11 mice/group).

##### Experiment 2

EEG recording for a baseline BSEEG score (as in Experiment 1) ran for 3 days. On day 5, we administered saline or LPS (0.5 mg/kg and 1.0 mg/kg) to young mice by i.p. injection, and sacrifices were conducted 6 hours after injection to assess the level of microglia activation by immunofluorescence (n□=□11 mice/group).

##### Experiment 3

On day -14, C57BL/6J aged mice (18-19 months old) underwent an EEG head mount implantation. The postoperative recovery period was 14 days. On day 1, we started EEG recording to measure the baseline BSEEG score for three days (days 1-3). On day 4, we conducted a battery of three types of behavioral tests (BFT, OFT, and Y-maze, as previously described). On day 5, we administered saline or LPS (0.1 mg/kg and 0.2 mg/kg) into aged mice by i.p. injection, and behavioral tests were repeated 6 hours after injection (n□=□11 mice/group). The reduced LPS doses for aged mice (0.1 mg/kg and 0.2 mg/kg) were determined because aged mice had higher rates of mortality at the same LPS doses used for young mice (0.5 mg/kg and 1.0 mg/kg) in a preliminary experiment (data not shown).

#### EEG electrode head mount implantation surgery

EEG electrode head mount implantation surgery was conducted as documented in previous studies^19,24^ (eMethods).

#### Behavioral tests

The battery of behavioral tests was performed as described in previous studies with modifications^17,18,25^. Each test was performed 24 hours before and 6 hours after a single injection of saline or LPS. Behavioral test results were calculated as a percentage of the baseline. The correlation between behavioral test endpoints and BSEEG score changes after LPS administration was examined. Mouse movement parameters were recorded via video camera and counted either manually by experimenters or automatically by the SMART 3.0 tracking system software (Panlab Harvard Apparatus, Holliston, MA; Cat# 76-0695).

##### Buried Food Test

BFT was implemented to specifically evaluate attention, which is typically affected in delirium. Although it is originally used to test the olfaction of the mice^26,27^, this method is also widely used in rodent animal studies for delirium to quantify attention and organized thinking^18,28,29^. The test was performed with modifications as described in previous studies^17,18,27^. Namely, 24 hours before the test, we removed all chow pellets from the cage environment but did not remove the water bottle (water ad libitum). The test cage was prepared with clean bedding that was 3 centimeters high. We buried one pellet 0.5 centimeters below the surface of the bedding so that it was not visible. We used around 2 g of the same chow pellet the animals were regularly fed. The food pellet was buried in a new and random location each time. We measured latency time, or the time from the mouse’s placement at the center of the test cage to the food pellet when it grasped the pellet in its forepaws and teeth. The mouse was allowed to consume the discovered pellet and was then returned to its home cage. The maximum observation time was set at 10 minutes in accordance with the literature^27^. If the mouse could not find the pellet within 10 minutes, the testing session ended, and the latency was defined as 600 seconds for that mouse. A new cage was used for each test to prevent the transmission of olfactory cues. We also changed gloves between tests.

##### Open Field Test

The open field test was performed to assess the mice’s locomotor activity and motor function^17,18,30,31^. The mouse was gently placed in center of an open field (40□×□40□×□40 cm (width□×□length□×□height), Stoelting, Wood Dale, IL; Cat# 10-000-294) and allowed to move freely for 5 minutes^25^. The total distance moved (meters), the time spent in center of the open field (seconds), the freezing time (seconds), and the latency to the first attempt in the center of the open field (seconds) were recorded and analyzed. The floor of the open field was cleaned with 70% ethanol solution between each test.

##### Y maze test

The Y maze consisted of three arms with an angle of 120° between adjacent arms (5□×□35□×□10 cm, Panlab Harvard Apparatus, Holliston, MA; Cat# 76-0079). The Y maze test is meant to quantify spatial and working memory based on spontaneous behavior without the need for reward or punishment. It can be performed quickly, as it only requires a single learning phase^32^. The test has been used in many studies that evaluate cognitive function in LPS-injected mice^30,33^. The Y maze test was performed as described in previous studies with modifications^17,18^. It consisted of 2 trials separated by an inter-trial interval (ITI). During the 10 minutes of the first trial (training), the mouse was allowed to explore two arms (the start arm and the other arm) of the maze, and the novel arm was blocked. After a 2-hour ITI, the second trial (retention) was conducted. For the second trial, the mouse was placed back in the start arm of the maze with free access to all three arms for 5□minutes. The duration in novel arm indicated the mouse’s level of spatial recognition memory. The entries in the novel arm indicated the spatial recognition memory and locomotor activity. The Y maze was cleaned with 70% ethanol between trials.

#### Immunofluorescence staining

Mouse brain slices were prepared for immunostaining to assess the effects of LPS-induced systemic inflammation on microglia in the hippocampal dentate gyrus (DG) and cortex regions. While mice were deeply anesthetized, perfusion with PBS (dilution 1:10; PBS (10X), pH 7.4, Gibco-Thermo Fisher Scientific, Waltham, MA; Cat# 70011044) and fixation with 4% paraformaldehyde (PFA) (dilution 1:4; Paraformaldehyde 16% Aqueous Solution EM Grade, Electron Microscopy Sciences, Hatfield, PA; Cat# 15710) were performed 6 hours after saline or LPS injection. The brain was dissected, and post-fixation was done overnight in 4% PFA at 4 °C. After fixation, the brain was dehydrated and preserved in 30% sucrose at 4 °C. Coronal frozen sections (40 μm thick) of the hippocampus and cortex were prepared by microtome (HM 450 Fully Automated Sliding Microtome, Epredia, Portsmouth, NH, USA). Brain slices were permeabilized with 1% Triton X-100 in PBS for 15 min and then blocked in 5% bovine serum albumin (BSA) (Sigma-Aldrich, St. Louis, MO; Cat# A3733) in PBS for 1 hour. To identify activated microglia, slices were incubated overnight at 4 °C in a blocking buffer containing an antibody against a cluster of differentiation (CD) 68 Rabbit (dilution 1:400; Cell Signaling Technology, Inc., Danvers, MA, USA; Cat# 97778)^34^. Slices were then incubated with fluorochrome-conjugated goat anti-rabbit IgG H&L secondary antibody (dilution 1:500; Alexa Fluor® 594, Invitrogen, Waltham, MA, USA; Cat# A32740) at room temperature for 2 hours. Next, slices were incubated for three days at 4 °C in a blocking buffer containing an antibody against ionized calcium-binding adaptor molecule 1 protein (IBA1) Goat (dilution 1:250; FUJIFILM Wako Pure Chemical Corporation, Osaka, Japan; Cat# 011-27991). Slices were subsequently incubated with donkey anti-goat-IgG antibody (dilution 1:500; Alexa Fluor® 488, Invitrogen; Cat# A11055) at room temperature for 2 hours. Finally, slices were counterstained with 4′, 6-diamidino-2-phenylindole (DAPI) (Vector Laboratories, Newark, CA; Cat# H-1800) at room temperature and observed by fluorescence microscopy (All-in-one fluorescence microscope BZ-X800, KEYENCE, Osaka, Japan). Microglia were quantified by a BZ-X800 analyzer that automatically counted the total number of IBA1 positive and CD68 positive cells in the hippocampal DG region at 10× magnification and in the cortex region at 4× magnification. Four slices were analyzed per mouse (n□=□11 mice/group) (2 (right and left) images/slice). Results are presented as fold change from the mean of the saline group. Results of IBA1 and CD68 images in the hippocampal DG and cortex are adjusted for size. Thus, scale bars are presented.

### Calculation of BSEEG score

#### EEG signal processing for BSEEG scores

Raw EEG recordings were digitally converted into power spectral density, following processing similar to that summarized in our previous clinical studies^9^, and then analyzed with an algorithm similar to that used in our human study. Sirenia^®^ software (Pinnacle Technology, Inc., Lawrence, KS) was used to record and export raw data. BSEEG scores were generated by calculating the ratio between 3 Hz power to 10 Hz power.

#### Web-based BSEEG score calculator

Raw EEG was digitally converted into power spectral density. We used the Sirenia software to record and export raw data into a European Data Format (EDF) file, which we processed through our web-based tool (https://sleephr.jp/gen/mice7days/). Our web-based tool automatically calculates BSEEG scores, the ratio of 3 Hz to 10 Hz power.

#### *Δ* BSEEG score

“Δ BSEEG score” was obtained from EEG2 (frontal electrode) to quantify the change in mouse BSEEG score after LPS injection. “Δ BSEEG score” was calculated from the mean BSEEG score over a 6-hour period after injection on Day 5, subtracted from the baseline BSEEG score. The baseline BSEEG score was calculated as the average BSEEG score for a 6-hour daytime period (12:00∼18:00) on days 1-3 (eFigure 2) and was set as zero.

#### Standardized BSEEG (sBSEEG) score

To highlight the change in BSEEG score from baseline in the 12-hour mean BSEEG score interval, the standardized BSEEG (sBSEEG) score was calculated as in our previous studies^19,24^, with a modification of the definition of the baseline BSEEG score. In this study, the sBSEEG score was the 12-hour mean BSEEG score subtracted from the baseline BSEEG score.

### Statistics

In all bar graphs, error bars were shown as a mean ± standard error of the mean (SEM). Normality was assessed using the Shapiro-Wilk normality test. Differences in variances of normally distributed data were assessed using Bartlett’s test. The differences between groups were analyzed with either one-way analysis of variance (ANOVA) followed by Bonferroni correction for multiple comparisons, where data were normally distributed and had the same variance across groups, or with the Kruskal-Wallis test followed by Dunn’s multiple comparison tests, where data were not normally distributed, or different variance across groups. A P value less than 0.05 was marked with an asterisk and considered statistically significant, and P values are indicated as follows in figures: ****P < 0.0001, ***P < 0.001, **P < 0.01, *P < 0.05. Correlations between normally distributed data were calculated using Pearson’s correlation, and between those that were not normally distributed were calculated using Spearman’s test. GraphPad Prism version 10 was used to analyze the data and create graphs.

#### Composite Z score

Composite Z score was derived from a battery of behavioral tests and reported as a scale to quantify the severity of postoperative impairments in mice consistent with the Confusion Assessment Method (CAM) feature in delirium^17^. The six behavioral tests of the composite Z score are as follows: 1) latency to eat food, 2) time spent in center, 3) latency to center, 4) freezing time, 5) entries in the novel arm, and 6) duration in novel arm. The Z score of each behavioral parameter was calculated as Z = [*ΔX _LPS_* - MEAN *(ΔX _Saline_)*]/SD *(ΔX _Saline_)*^25^. In the formula, *ΔX _Saline_* was the difference between the score of mice in the saline group at 6 hours after injection and the baseline score. *ΔX _LPS_*was the difference between the score of mice in the LPS (0.5 mg/kg or 1.0 mg/kg) group at 6 hours after injection and the score at baseline; MEAN *(ΔX _Saline_)* was the mean of *ΔX _Saline_*, and SD *(ΔX _Saline_)* was the standard deviation of *ΔX _Saline_*. Composite Z score was calculated as the sum of 6 Z scores normalized with the SD calculated from the sum of Z scores in the saline group. Given that the reduction in time spent in center, the freezing time (open field test), the reduction in duration, and entries in the novel arm (Y maze test) indicate impairment of the movement, we multiplied the Z score values representing these behaviors by -1 before calculating the composite Z score.

## Results

### The relationship between BSEEG score and behavior change after LPS injection in young mice (Experiment 1)

The typical pattern of sBSEEG score changes and the effect of LPS injection was reproducible as in previous studies. There were regular diurnal changes in every 12-hour sBSEEG score. sBSEEG scores increased, and regular diurnal changes were disrupted after LPS injection (Figure 1A). There were significant differences between saline and LPS 0.5 mg/kg and 1.0 mg/kg groups in Δ BSEEG score (Figure 1B). There were significant differences between saline and LPS 0.5 mg/kg and 1.0 mg/kg groups in total distance in OFT (Figure 1C). There was a moderate negative correlation between Δ BSEEG score and total distance in OFT (Figure 1D). There was a significant difference between saline and LPS 1.0 mg/kg groups in latency time in BFT (Figure 1E). There was a strong positive correlation between Δ BSEEG score and latency time in BFT (Figure 1F). There were no significant differences in time spent in novel arm in the Y maze test (Figure 1G). There was a weak negative correlation between Δ BSEEG score and duration in novel arm in the Y maze test (Figure 1H). There was significant difference between saline and LPS 1.0 mg/kg groups in entries in the Y maze test (Figure 1I). There was a moderate negative correlation between Δ BSEEG score and entries in Y maze test (Figure 1J). There were significant differences between saline and LPS 0.5 mg/kg and 1.0 mg/kg groups in composite Z score (Figure 1K and eTable 2). There was a moderate positive correlation between Δ BSEEG score and composite Z score (Figure 1L). We also assessed whether latency to the pellet in BFT was caused by low locomotor activity. Latency to pellet in BFT was multiplied by total distance in OFT to adjust the influence of hypoactivity. The correlation between latency to pellet in BFT and Δ BSEEG score remained strong even after adjustment for hypoactivity (eFigure 3A). Similarly, we divided the duration in novel arm in the Y maze test by total distance in OFT. There was no correlation between the duration in novel arm in the Y maze test and Δ BSEEG score (eFigure 3B).

**Figure 1.**
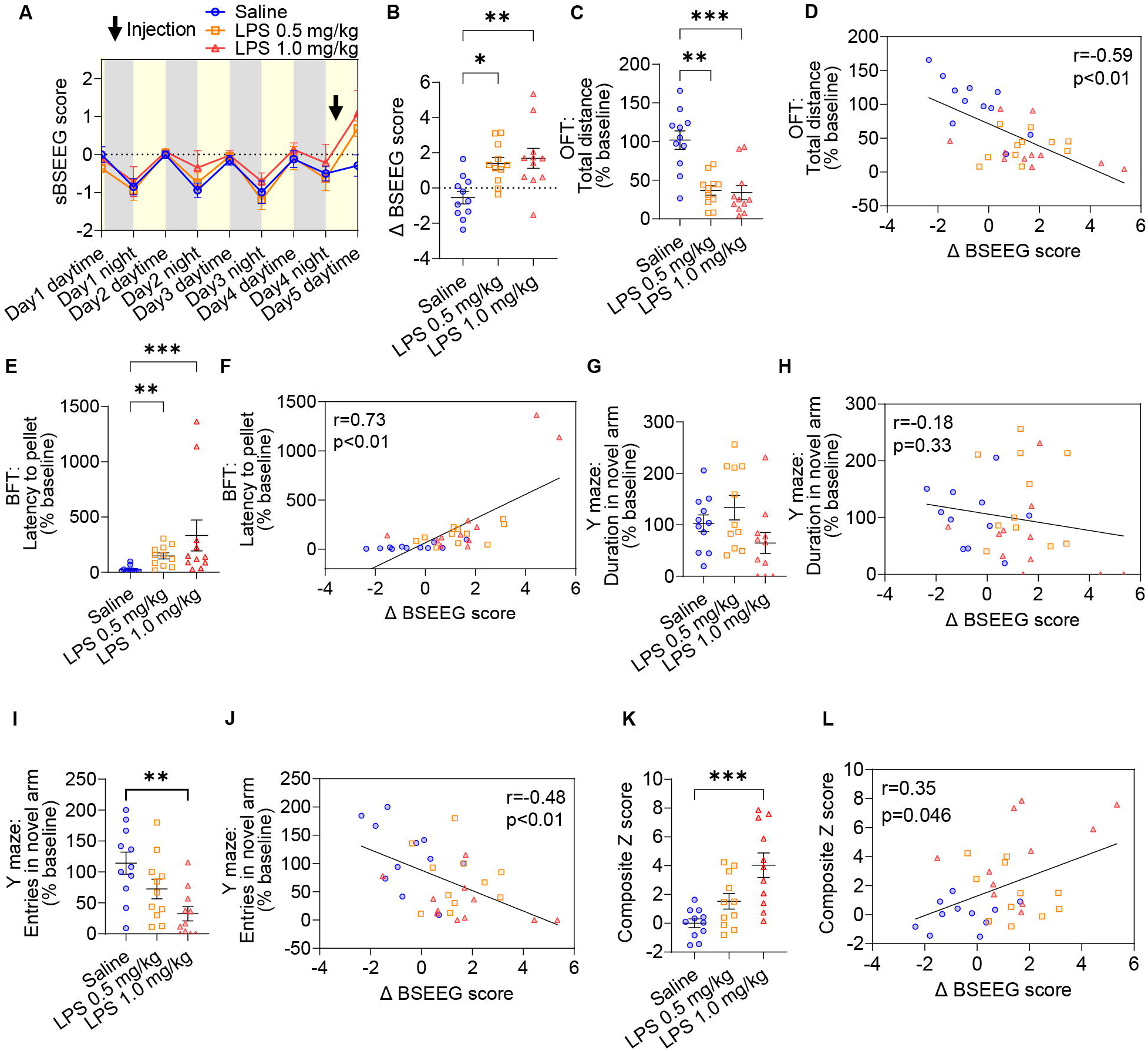
The relationship between BSEEG score and LPS-induced behavior in young mice (Experiment 1) (Saline, n=11; LPS 0.5 mg/kg, n=11; LPS 1.0 mg/kg, n=11). (A) A time course of 12-hour average sBSEEG scores after saline/LPS injection in young mice. (B) Comparison of Δ BSEEG scores on Day 5 between saline and LPS 0.5 mg/kg and 1.0 mg/kg groups (Saline vs. LPS 0.5 mg/kg: p = 0.012, Saline vs. LPS 1.0 mg/kg: p = 0.0032; mean: Saline = -0.55, LPS 0.5 mg/kg = 1.39, LPS 1.0 mg/kg = 1.69). (C) Comparison of total distance in OFT between saline and LPS 0.5 mg/kg and 1.0 mg/kg groups (Saline vs. LPS 0.5 mg/kg: p = 0.0052, Saline vs. LPS 1.0 mg/kg: p = 0.0005; mean: Saline = 102%, LPS 0.5 mg/kg = 37%, LPS 1.0 mg/kg = 34%). (D) Correlation between Δ BSEEG score and total distance in OFT. (E) Comparison of latency to pellet in BFT between saline and LPS 0.5 mg/kg and 1.0 mg/kg groups (Saline vs. LPS 0.5 mg/kg: p = 0.0022, Saline vs. LPS 1.0 mg/kg: p = 0.0008; mean: Saline = 25%, LPS 0.5 mg/kg = 148%, LPS 1.0 mg/kg = 333%). (F) Correlation between Δ BSEEG score and latency to pellet in BFT. (G) Comparison of duration in novel arm in Y maze test between saline and LPS 0.5 mg/kg and 1.0 mg/kg groups (mean: Saline = 103%, LPS 0.5 mg/kg = 133%, LPS 1.0 mg/kg = 65%). (H) Correlation between Δ BSEEG score and time in novel arm in Y maze test. (I) Comparison of entries in novel arm in Y maze test between saline and LPS 0.5 mg/kg and 1.0 mg/kg groups (Saline vs. LPS 1.0 mg/kg: p = 0.0029; mean: Saline = 114%, LPS 0.5 mg/kg = 72%, LPS 1.0 mg/kg = 32%). (J) Correlation between Δ BSEEG score and entries in novel arm in Y maze test. (K) Comparison of composite Z score between saline and LPS 0.5 mg/kg and 1.0 mg/kg groups (Saline vs. LPS 1.0 mg/kg: p = 0.0009; mean: Saline = 0, LPS 0.5 mg/kg = 1.53, LPS 1.0 mg/kg = 4.03). (L) Correlation between Δ BSEEG score and composite Z score.

### The relationship between BSEEG score and microglia activation after LPS injection in young mice (Experiment 2)

We investigated the relationship between BSEEG score and microglia activation in the hippocampal DG. As shown previously, LPS injection increased the BSEEG score in the present study (Figure 2A). There were significant differences between saline and LPS 0.5 mg/kg and 1.0 mg/kg group in IBA1 positive cells in the hippocampal DG region (Figure 2B and 2C) (eFigure 4A). The positive correlation between Δ BSEEG score and IBA1 positive cells in the hippocampal DG region was strong (Figure 2D). There were significant differences between saline and LPS 0.5 mg/kg and 1.0 mg/kg group in CD68 positive cells in hippocampal DG region (Figure 2B and 2E) (eFigure 4B). The positive correlation between Δ BSEEG score and CD68 positive cells in the hippocampal DG region was strong (Figure 2F). We also investigated the relationship between the BSEEG score and microglia activation in the cortex. There were significant differences between IBA1 positive cells in the cortex among saline and LPS 0.5 mg/kg and 1.0 mg/kg group (Figure 3A and 3B) (eFigure 4C). The positive correlation between Δ BSEEG score and IBA1 positive cells in the cortex was strong (Figure 3C). There were significant differences between saline and LPS 0.5 mg/kg and 1.0 mg/kg group in CD68 positive cells in the cortex (Figure 3A and 3D) (eFigure 4D). The positive correlation between Δ BSEEG score and CD68 positive cells in the cortex was strong (Figure 3E).

**Figure 2.**
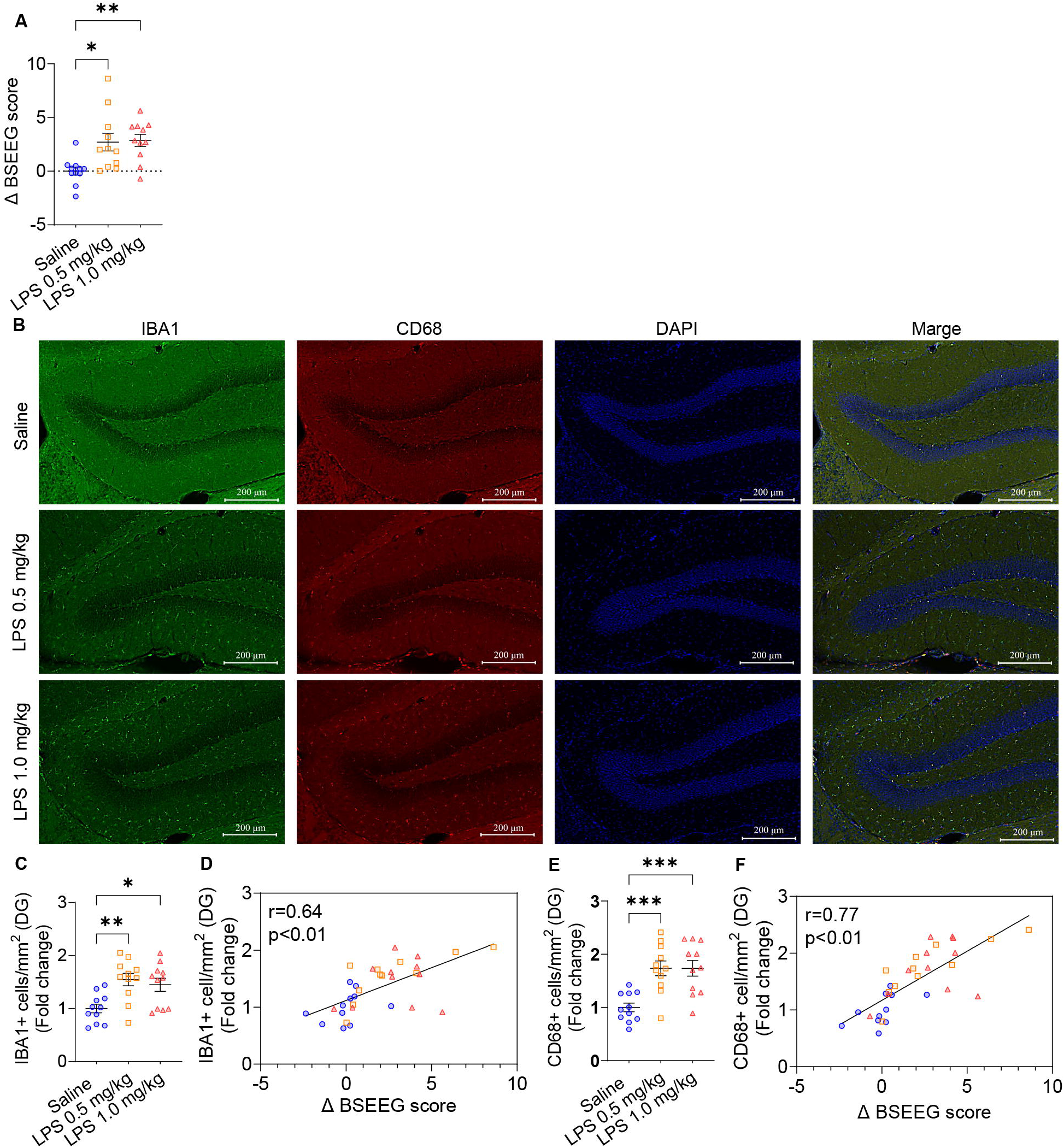
The relationship between the BSEEG score and IBA1 and CD68 positive cell count in hippocampal DG 6 hours after LPS i.p. injection in young mice (Experiment 2) (Saline, n=11; LPS 0.5 mg/kg, n=11; LPS 1.0 mg/kg, n=11). (A) Comparison of Δ BSEEG score between saline and LPS 0.5 mg/kg and 1.0 mg/kg group (Saline vs. LPS 0.5 mg/kg: p = 0.013, Saline vs. LPS 1.0 mg/kg: p = 0.0078; mean: Saline = 0.01, LPS 0.5 mg/kg = 2.70, LPS 1.0 mg/kg = 2.87). (B) Immunofluorescence images of hippocampal DG (Scale bar = 200 µm). (C) IBA1 positive cell count per mouse in hippocampal DG (Saline vs. LPS 0.5 mg/kg: p = 0.004, Saline vs. LPS 1.0 mg/kg: p = 0.021; mean: Saline = 1.00, LPS 0.5 mg/kg = 1.55, LPS 1.0 mg/kg = 1.45). (D) The relationship between BSEEG score and IBA1 positive cell count per mouse in hippocampal DG. (E) CD68 positive cell count per mouse in hippocampal DG (Saline vs. LPS 0.5 mg/kg: p = 0.0008, Saline vs. LPS 1.0 mg/kg: p = 0.0008; mean: Saline = 1.00, LPS 0.5 mg/kg = 1.74, LPS 1.0 mg/kg = 1.74). (F) The relationship between BSEEG score and CD68 positive cell count per mouse in hippocampal DG.

**Figure 3.**
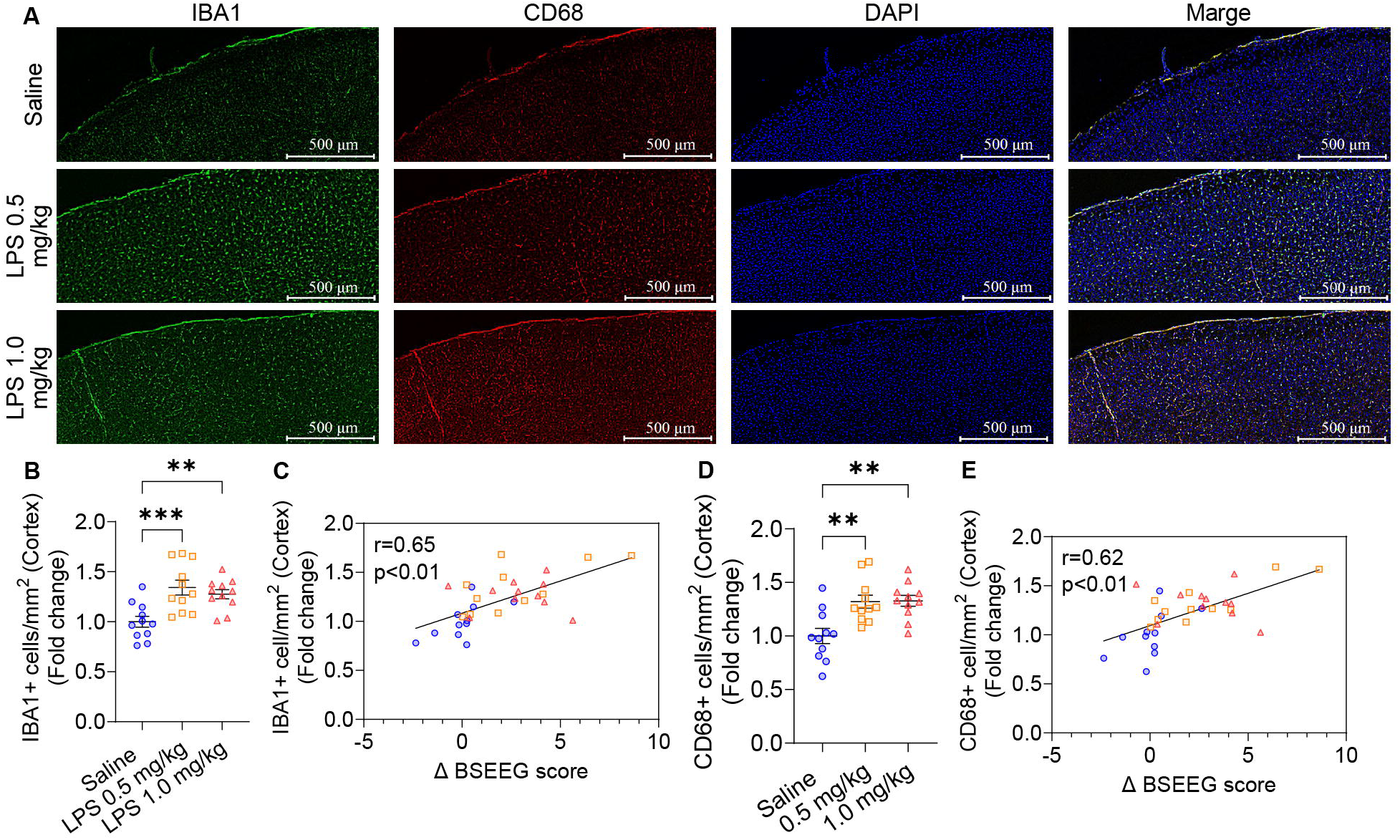
The relationship between the BSEEG score and IBA1 and CD68 positive cell count in cortex 6 hours after LPS i.p. injection in young mice (Experiment 2) (Saline, n=11; LPS 0.5 mg/kg, n=11; LPS 1.0 mg/kg, n=11). (A) Immunofluorescence images of the cortex (Scale bar = 500 µm). (B) IBA1 positive cell count per mouse in cortex (Saline vs. LPS 0.5 mg/kg: p = 0.0009, Saline vs. LPS 1.0 mg/kg: p = 0.0076; mean: Saline = 1.00, LPS 0.5 mg/kg = 1.34, LPS 1.0 mg/kg = 1.28). (C) The relationship between BSEEG score and IBA1 positive cell count per mouse in cortex. (D) CD68 positive cell count per mouse in cortex (Saline vs. LPS 0.5 mg/kg: p = 0.0028, Saline vs. LPS 1.0 mg/kg: p = 0.0022; mean: Saline = 1.00, LPS 0.5 mg/kg = 1.320, LPS 1.0 mg/kg = 1.328). (E) The relationship between BSEEG score and CD68 positive cell count per mouse in cortex.

### The relationship between BSEEG score and behavior change and microglia activation after LPS injection in aged mice (Experiment 3)

Next, we assessed the relationship between BSEEG score and behavior change as well as microglia activation after LPS injection in aged mice. The same experiment schedule as in Experiments 1 and 2 was used. However, there was a reduced LPS dosage (0.1 mg/kg and 0.2 mg/kg) than that given to younger mice. Unlike young mice who showed regular sBSEEG score diurnal changes, aged mice did not show such regular sBSEEG score diurnal changes (Figure 4A). We observed a similar increase in BSEEG score following LPS injection (Figure 4A). There were significant differences between saline and LPS 0.1 mg/kg and 0.2 mg/kg group in Δ BSEEG score (Figure 4B). Regarding the behavioral test, there was a significant decrease in total distance in OFT between the saline and LPS 0.1 mg/kg group (Figure 4C). The negative correlation between Δ BSEEG score and total distance in OFT was strong (Figure 4D). Activity levels were too low to assess other behavioral tests. Almost all mice failed to find pellets in BFT or enter novel arm in the Y maze test.

**Figure 4.**
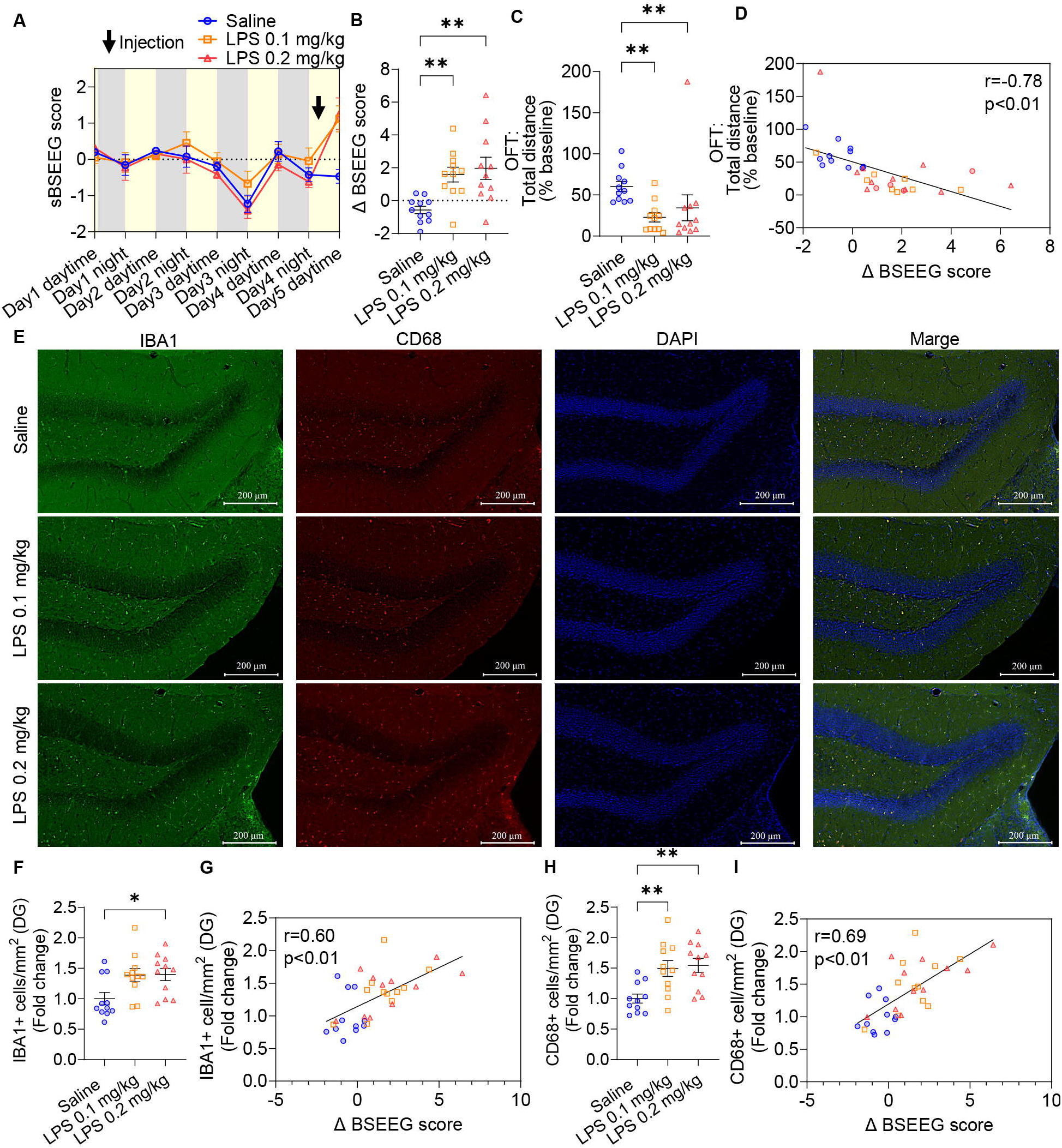
The relationship between BSEEG score and IBA1 and CD68 positive cell count in hippocampal DG 6 hours after LPS i.p. injection in aged mice (Experiment 3) (Saline, n=11; LPS 0.1 mg/kg, n=11; LPS 0.2 mg/kg, n=11). (A) A time course of 12-hour average sBSEEG scores after saline/LPS injection in aged mice. (B) Comparison of Δ BSEEG scores between saline and LPS 0.1 mg/kg and 0.2 mg/kg group (Saline vs. LPS 0.1 mg/kg: p = 0.0031, Saline vs. LPS 0.2 mg/kg: p = 0.0033; mean: Saline = -0.57, LPS 0.1 mg/kg = 1.59, LPS 0.2 mg/kg = 1.97). (C) Comparison of total distance in OFT between saline and LPS 0.1 mg/kg and 0.2 mg/kg group (Saline vs. LPS 0.1 mg/kg: p = 0.0024, Saline vs. LPS 0.2 mg/kg: p = 0.0042; mean: Saline = 60.28, LPS 0.1 mg/kg = 22.68, LPS 0.2 mg/kg = 34.32). (D) Correlation between Δ BSEEG score and total distance in OFT. (E) Immunofluorescence images in hippocampal DG (Scale bar = 200 µm). (F) IBA1 positive cell count (Saline vs. LPS 0.2 mg/kg: p = 0.015; mean: Saline = 1.00, LPS 0.1 mg/kg = 1.39, LPS 0.2 mg/kg = 1.40). (G) The relationship between BSEEG score and IBA1 positive cell count. (H) CD68 positive cell count (Saline vs. LPS 0.1 mg/kg: p = 0.0095, Saline vs. LPS 0.2 mg/ kg: p = 0.004; mean: Saline = 1.00, LPS 0.1 mg/kg = 1.49, LPS 0.2 mg/kg = 1.55). (I) The relationship between BSEEG score and CD68 positive cell count.

Also, in aged mice, we investigated the relationship between the BSEEG score and microglia activation in the hippocampal DG region. There were significant differences between saline and LPS 0.1 mg/kg and 0.2 mg/kg group in IBA1 positive cells in the hippocampal DG region (Figure 4E and 4F) (eFigure 5A). The positive correlation between Δ BSEEG score and IBA1 positive cells in the hippocampal DG region was strong (Figure 4G). There were significant differences between saline and LPS 0.1 mg/kg and 0.2 mg/kg group in CD68 positive cells in hippocampal DG region (Figure 4E and 4H) (eFigure 5B). The positive correlation between Δ BSEEG score and CD68 positive cells in the hippocampal DG region was strong (Figure 4I).

Moreover, we investigated the relationship between the BSEEG score and microglia activation in the cortex in aged mice. There were significant differences between saline and LPS 0.1 mg/kg and 0.2 mg/kg group in IBA1 positive cells in the cortex (Figure 5A and 5B) (eFigure 5C). The positive correlation between Δ BSEEG score and IBA1 positive cells in the cortex was strong (Figure 5C). There were significant differences between saline and LPS 0.1 mg/kg and 0.2 mg/kg group in CD68 positive cells in the cortex (Figure 5A and 5D) (eFigure 5D). The positive correlation between Δ BSEEG score and CD68 positive cells in the cortex was strong (Figure 5E).

**Figure 5.**
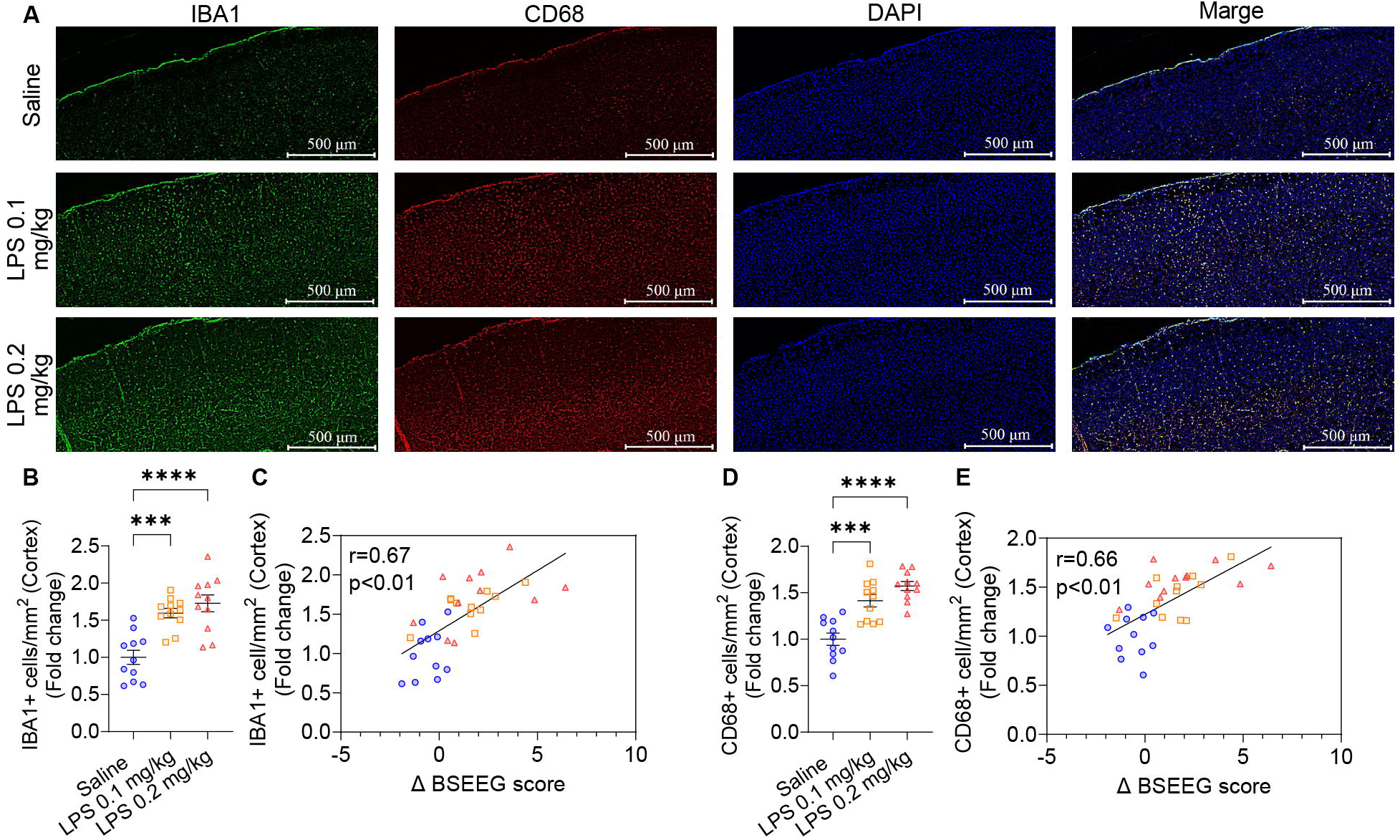
The relationship between BSEEG score and IBA1 and CD68 positive cell count in cortex 6 hours after LPS i.p. injection in aged mice (Experiment 3) (Saline, n=11; LPS 0.1 mg/kg, n=11; LPS 0.2 mg/kg, n=11). (A) Immunofluorescence images in cortex (Scale bar = 500 µm). (B) IBA1 positive cell count per mouse (Saline vs. LPS 0.1 mg/kg: p = 0.0003, Saline vs. LPS 0.2 mg/kg: p < 0.0001; mean: Saline = 1.00, LPS 0.1 mg/kg = 1.60, LPS 0.2 mg/kg = 1.73). (C) The relationship between BSEEG score and IBA1 positive cell count per mouse. (D) CD68 positive cell count per mouse (Saline vs. LPS 0.1 mg/kg: p = 0.0001, Saline vs. LPS 0.2 mg/kg: p < 0.0001; mean: Saline = 1.00, LPS 0.1 mg/kg = 1.41, LPS 0.2 mg/kg = 1.57). (E) The relationship between BSEEG score and CD68 positive cell count per mouse.

## Discussion

This study aimed to verify whether an increase in BSEEG scores induced by LPS in mice correlates with or could even be an alternative indicator for delirium-like behavior or microglia activation.

Several papers use a behavioral test battery consisting of BFT, OFT, and Y maze test to evaluate postoperative delirium-like behavior^17,18^. The behavioral test battery has also been used to assess delirium-like behavior following LPS administration^35^. Therefore, in experiment 1, we conducted EEG recordings and the behavioral test battery to evaluate delirium-like behavior after LPS administration. We also verified the correlation between BSEEG scores and delirium-like behavior after LPS administration. LPS administration led to an increase in BSEEG scores and, in the behavioral test battery, a decrease in locomotor activity, an increase in attention deficits, and an increase in the composite Z-score, although no spatial cognitive impairment was observed. Remarkably, BSEEG scores correlated with decreased locomotor activity, attention deficits, and composite Z-score. In particular, the correlation between BSEEG scores and the composite Z-score, which is used to evaluate the severity of postoperative delirium, suggests that the BSEEG score could potentially indicate the severity of delirium-like behavior as an alternative to behavioral tests.

In experiment 2, we evaluated the correlation between BSEEG scores and microglia activation in the dentate gyrus and cerebral cortex regions through immunofluorescence staining with both IBA1, a marker for microglia and macrophages, and CD68, a marker for activated microglia, after LPS administration. It has been previously reported that the hippocampus is a key region affected by LPS exposure^36^. The hippocampal dentate gyrus is known to be the area where neurogenesis occurs in the adult rodent brain^37^. It has been reported that a single i.p. injection of LPS induced significant microglial activation and acute neuroinflammation in the hippocampal DG region^38^. Thus, we investigated the relationship between BSEEG score and microglia activation in the hippocampal DG region to verify our hypothesis that the BSEEG score reflects the severity of neuroinflammation in delirium. Also, EEG primarily captures electric signals from regions of the brain close to the surface, such as the cortex, while BSEEG specifically captures slowing in brain wave activity, which theoretically reflects pathological phenomena in the cortex. For this reason, we assessed microglial activation in the cortex as well. In response to LPS administration, BSEEG scores increased as expected. There was also an increase in the number of IBA1 and CD68 positive cells in both the dentate gyrus and cerebral cortex regions after LPS administration. These results are consistent with previous studies reporting that LPS administration induces systemic inflammation and increases the number of activated microglia^22,39^. Furthermore, there were positive correlations between BSEEG scores and the number of IBA1 and CD68 positive cells. We believe that these findings support our hypothesis that an increase in BSEEG scores is attributable to neuroinflammation.

Delirium often occurs in older patients, and age is known to be a risk factor for delirium^40^. Therefore, in experiment 3, we evaluated delirium-like behavior and microglia activation after LPS administration in aged mice. However, LPS administration significantly reduced locomotor activity in the aged mice, making behavioral testing unfeasible. LPS administration in aged mice also increased BSEEG scores and the number of IBA1 and CD68 positive cells in both the dentate gyrus and cerebral cortex regions. Furthermore, there was a positive correlation between BSEEG scores and the number of IBA1 and CD68 positive cells. Consequently, we suggest that even when behavioral tests are not feasible due to reduced locomotor activity (hypoactivity) from LPS administration, the BSEEG method can detect delirium-like states accompanied by microglia activation from LPS administration. These findings suggest that the BSEEG method could be useful for detecting LPS-induced delirium-like states, including hypoactive ones, in mice of any age. On the other hand, the increase in BSEEG score may also be able to capture other hypoactive states which are different from delirium, such as non-delirious encephalopathy, sepsis, or drug toxicity. In fact, our previous clinical study reported that there might also be an association between the BSEEG score and sepsis^41^. It may be interest to investigate the specificity of the BSEEG method with not only delirium buy also with non-delirious hypoactivity states. We recognize that there are several limitations to this study. First, our behavioral battery was conducted only once instead of multiple times a day as in previous literature. This might not capture the one of the core features in delirium, the fluctuating nature of the behavioral changes Also, there might be better behaviors in aged mice at later time points or with lower dose of LPS. These trials are worth trying in the future. However, it is reported that LPS suppressed the locomotor activity of even young mice for several days^42^ and did that of aged mice at least for 24hours^43^. On the other hand, in a previous study, the peaks of the BSEEG increase were at post 6 hours or 12 hours LPS injection and the increase in BSEEG scores with lower dose of LPS might be expected to be insufficient due to the feature of dose-dependent BSEEG score increase in response to LPS^19^. Thus, to achieve the goal to assess the correlation between the peak of BSEEG score and behavioral tests as well as microglial activation, we focused on the first 6 hours peaks of the increase in BSEEG scores after sufficient dose LPS injection in this study. Second, we did not directly measure inflammatory cytokines in the brain. Some studies suggest that activated microglia do not inherently determine their function by themselves as either pro- or anti-inflammatory, but rather, a balance of inflammatory or anti-inflammatory secretory factors influence their ultimate function^44,45^. However, LPS administration is known to induce systemic inflammation and activate pro-inflammatory microglia^46^. Thus, our data confirming the correlation between BSEEG scores and microglia activation is confirmatory of our hypothesis without necessarily measuring inflammatory cytokines. Third, the causal relationships that might underlie the correlations that we have identified remain uncertain. This ambiguity needs to be addressed in future studies. Fourth, it is difficult to distinguish delirium-like behavior from sickness behavior in mice to whom LPS is administered. LPS is known to induce lethargy, depression-like behavior, and reduced activity in animals, decreasing their motivation to move such that energy is conserved for metabolism. These are all features of sickness behavior^47^. Also, as one of the sickness behaviors, even low dose LSP injection reduces food intake^48^. Moreover, it is reported that there are olfaction declines in aged mice^49^ and with LPS^50^. Thus, there might be some confounders in behavioral tests, such as the effect of olfaction, the reduction of food intake, and the reduction in locomotor activity. To mitigate the impact of sickness behavior on locomotor activity, we adjusted the latency to the pellet in BFT and the time spent in novel arm in Y-maze test by locomotor activity. Even after adjustment, however, attention deficits remained correlated with BSEEG scores. This suggests that BSEEG scores may reflect attention deficits in delirium-like behaviors. To verify this suggestion, we need to investigate the relation between BSEEG scores and other delirium models, having less confounders, like a surgery model in the future. Fifth, the reduction in locomotor activity induced by LPS administration might have had broader effects on other tested parameters. In this study, we aimed to evaluate the severity of delirium by calculating a composite Z-score equivalent to CAM using the BFT, OFT, and Y-maze behavioral test battery. Since the composite Z-score is calculated from multiple parameters in each behavioral test, locomotor activity differences may have influenced the composite Z-score of each group. However, the formula for composite Z-score makes it impossible to exclude the influence of differences in locomotor activity. Sixth, our study included only male mice. The reason is that including females complicates age comparisons due to changing female hormone levels with age. To overcome this limitation, we need a study including female mice in the future.

In conclusion, by administering LPS to mice and evaluating delirium-like states, we succeeded in reaffirming the validity of the BSEEG method at both behavioral and cellular levels. We confirmed that the BSEEG method correlates with reduced locomotor activity, attention deficits, severity of delirium-like states, and microglia activation induced by LPS administration. Furthermore, the BSEEG method may potentially measure delirium-like states that are difficult to evaluate through behavioral tests in aged mice due to reduced locomotor activity caused by LPS administration. Thus, we suggest that the BSEEG method is useful for detecting LPS-induced delirium-like states in mice regardless of age. Future studies should investigate the utility of BSEEG in other delirium models, such as postoperative delirium models.

## Acknowledgments

No funding was received for conducting this study. We appreciate the support from the Stanford University School of Medicine Veterinary Service Center (VSC).

## Author contributions

GS conceptualized, designed, and coordinated the entire research study. TN designed and conducted experiments, acquired data, analyzed data, and wrote the initial draft of the manuscript. KY designed immunofluorescence staining and conducted perfusion and fixation. NG conducted behavioral tests, sliced brains, and analyzed data. AS programmed the web-based BSEEG score calculator. KY, NG, AS, TS, TI, BA, JRM, NJP, NJD, TY, and MI critically reviewed the manuscript. TN and GS wrote the final version of the manuscript.

## Appendixes (Supplemental Material titles and legends)

eMthods

eTable 1. All reagent or resource.

eTable 2. Z scores on all mice.

eFigure 1. Body weight after the head mount implantation surgery.

Experiment 1; Saline, n=11; LPS 0.5 mg/kg, n=11; LPS 1.0 mg/kg, n=11; Saline vs. LPS 0.5 mg/kg: p = 0.76, Saline vs. LPS 1.0 mg/kg: p = 0.33, LPS 0.5 mg/kg vs. LPS 1.0 mg/kg: p = 1.00; mean: Saline = 27, LPS 0.5 mg/kg = 25.91, LPS 1.0 mg/kg = 25.45.

Experiment 2; Saline, n=11; LPS 0.5 mg/kg, n=11; LPS 1.0 mg/kg, n=11; Saline vs. LPS 0.5 mg/kg: p = 1.00, Saline vs. LPS 1.0 mg/kg: p = 1.00, LPS 0.5 mg/kg vs. LPS 1.0 mg/kg: p = 1.00; mean: Saline = 25.82, LPS 0.5 mg/kg = 26.18, LPS 1.0 mg/kg = 26.09.

Experiment 3; Saline, n=11; LPS 0.1 mg/kg, n=11; LPS 0.2 mg/kg, n=11; Saline vs. LPS 0.1 mg/kg: p = 1.00, Saline vs. LPS 0.2 mg/kg: p = 1.00, LPS 0.1 mg/kg vs. LPS 0.2 mg/kg: p = 1.00; mean: Saline = 40, LPS 0.1 mg/kg = 39.64, LPS 0.2 mg/kg = 43.00.

eFigure 2. Experiment Schedules.

eFigure 3. The correlation between each parameter adjusted by the total distance in OFT with Δ BSEEG score (Experiment 1) (Saline, n=11; LPS 0.5 mg/kg, n=11; LPS 1.0 mg/kg, n=11). (A) The correlation between latency to pellet in BFT adjusted by total distance in OFT and Δ BSEEG score. (B) The correlation between duration in novel arm in Y maze test adjusted by total distance in OFT and Δ BSEEG score.

eFigure 4. Brain slice images (88/group) cell counts in the hippocampal DG and cortex 6 hours after LPS i.p. injection in young mice (Experiment 2) (Saline, n=11; LPS 0.5 mg/kg, n=11; LPS 1.0 mg/kg, n=11) (2 (right and left) images/slice) (4 slices/mouse).

(A) IBA1 positive cell counts in images of brain slices from hippocampal DG (Saline vs. LPS 0.5 mg/kg: p < 0.0001, Saline vs. LPS 1.0 mg/kg: p < 0.0001).

(B) CD68 positive cell counts in images of brain slices from hippocampal DG (Saline vs. LPS 0.5 mg/kg: p < 0.0001, Saline vs. LPS 1.0 mg/kg: p < 0.0001).

(C) IBA1 positive cell counts in images of brain slices from cortex (Saline vs. LPS 0.5 mg/kg: p < 0.0001, Saline vs. LPS 1.0 mg/kg: p < 0.0001).

(D) CD68 positive cell counts in images of brain slices from cortex (Saline vs. LPS 0.5 mg/kg: p < 0.0001, Saline vs. LPS 1.0mg/kg: p < 0.0001).

eFigure 5. Brain slice images (88/group) cell counts in the hippocampal DG and cortex 6 hours after LPS i.p. injection in aged mice (Experiment 3) (Saline, n=11; LPS 0.1 mg/kg, n=11; LPS 0.2 mg/kg, n=11) (2 (right and left) images/slice) (4 slices/mouse).

(A) IBA1 positive cell counts on brain slice images in the hippocampal DG (Saline vs. LPS 0.1 mg/kg: p < 0.0001, Saline vs. LPS 0.2 mg/kg: p < 0.0001).

(B) CD68 positive cell counts on brain slice images in the hippocampal DG (Saline vs. LPS 0.1 mg/kg: p < 0.0001, Saline vs. LPS 0.2 mg/kg: p < 0.0001).

(C) IBA1 positive cell counts on brain slice images in the cortex (Saline vs. LPS 0.1 mg/kg: p < 0.0001, Saline vs. LPS 0.2 mg/kg: p < 0.0001).

(D) CD68 positive cell count on brain slice images in the cortex (Saline vs. LPS 0.1 mg/kg: p < 0.0001, Saline vs. LPS 0.2 mg/kg: p < 0.0001, LPS 0.1 mg/kg vs. LPS 0.2 mg/kg: p = 0.0064).

## References

1. Wilson JE, Mart MF, Cunningham C, et al. Delirium. Nat Rev Dis Primers. Nov 12 2020;6(1):90. doi:10.1038/s41572-020-00223-4

2. Fong TG, Tulebaev SR, Inouye SK. Delirium in elderly adults: diagnosis, prevention and treatment. Nat Rev Neurol. Apr 2009;5(4):210–20. doi:10.1038/nrneurol.2009.24

3. Inouye SK. Delirium in older persons. N Engl J Med. Mar 16 2006;354(11):1157–65. doi:10.1056/NEJMra052321

4. Inouye SK, Westendorp RG, Saczynski JS. Delirium in elderly people. Lancet. Mar 08 2014;383(9920):911–22. doi:10.1016/S0140-6736(13)60688-1

5. Witlox J, Eurelings LS, de Jonghe JF, Kalisvaart KJ, Eikelenboom P, van Gool WA. Delirium in elderly patients and the risk of postdischarge mortality, institutionalization, and dementia: a meta-analysis. JAMA. Jul 28 2010;304(4):443–51. doi:10.1001/jama.2010.1013

6. Francis J, Young GB. Diagnosis of delirium and confusional states. UpToDate: Waltham, MA. 2012;

7. Inouye SK. The dilemma of delirium: clinical and research controversies regarding diagnosis and evaluation of delirium in hospitalized elderly medical patients. Am J Med. Sep 1994;97(3):278–88. doi:10.1016/0002-9343(94)90011-6

8. Setters B, Solberg LM. Delirium. Prim Care. Sep 2017;44(3):541–559. doi:10.1016/j.pop.2017.04.010

9. Shinozaki G, Chan AC, Sparr NA, et al. Delirium detection by a novel bispectral electroencephalography device in general hospital. Psychiatry Clin Neurosci. Dec 2018;72(12):856–863. doi:10.1111/pcn.12783

10. Nishizawa Y, Yamanashi T, Saito T, et al. Bispectral EEG (BSEEG) Algorithm Captures High Mortality Risk Among 1,077 Patients: Its Relationship to Delirium Motor Subtype. Am J Geriatr Psychiatry. Mar 08 2023;doi:10.1016/j.jagp.2023.03.002

11. Zarei K, Sparr NA, Trapp NT, et al. Bispectral EEG (BSEEG) to assess arousal after electro-convulsive therapy (ECT). Psychiatry Res. Jan 25 2020;285:112811. doi:10.1016/j.psychres.2020.112811

12. Lee S, Yuki K, Chan A, Cromwell J, Shinozaki G. The point-of-care EEG for delirium detection in the emergency department. Am J Emerg Med. May 2019;37(5):995–996. doi:10.1016/j.ajem.2018.10.004

13. Shinozaki G, Bormann NL, Chan AC, et al. Identification of high mortality risk patients and prediction of outcomes in delirium by Bispectral EEG. J Clin Psychiatry. 2019;80(5)doi:10.4088/JCP.19m12749

14. Yamanashi T, Crutchley KJ, Wahba NE, et al. Evaluation of point-of-care thumb-size bispectral electroencephalography device to quantify delirium severity and predict mortality. Br J Psychiatry. Aug 2 2021:1–8. doi:10.1192/bjp.2021.101

15. Dilger RN, Johnson RW. Aging, microglial cell priming, and the discordant central inflammatory response to signals from the peripheral immune system. J Leukoc Biol. Oct 2008;84(4):932–9. doi:10.1189/jlb.0208108

16. Schreuder L, Eggen BJ, Biber K, Schoemaker RG, Laman JD, de Rooij SE. Pathophysiological and behavioral effects of systemic inflammation in aged and diseased rodents with relevance to delirium: A systematic review. Brain Behav Immun. May 2017;62:362–381. doi:10.1016/j.bbi.2017.01.010

17. Peng M, Zhang C, Dong Y, et al. Battery of behavioral tests in mice to study postoperative delirium. Sci Rep. 07 20 2016;6:29874. doi:10.1038/srep29874

18. Illendula M, Osuru HP, Ferrarese B, et al. Surgery, Anesthesia and Intensive Care Environment Induce Delirium-Like Behaviors and Impairment of Synaptic Function-Related Gene Expression in Aged Mice. Front Aging Neurosci. 2020;12:542421. doi:10.3389/fnagi.2020.542421

19. Yamanashi T, Malicoat JR, Steffen KT, et al. Bispectral EEG (BSEEG) quantifying neuro-inflammation in mice induced by systemic inflammation: A potential mouse model of delirium. J Psychiatr Res. Jan 2021;133:205–211. doi:10.1016/j.jpsychires.2020.12.036

20. Kuswardhani RAT, Sugi YS. Factors Related to the Severity of Delirium in the Elderly Patients With Infection. Gerontol Geriatr Med. 2017;3:2333721417739188. doi:10.1177/2333721417739188

21. Esme M, Topeli A, Yavuz BB, Akova M. Infections in the Elderly Critically-Ill Patients. Front Med (Lausanne). 2019;6:118. doi:10.3389/fmed.2019.00118

22. Hoogland IC, Houbolt C, van Westerloo DJ, van Gool WA, van de Beek D. Systemic inflammation and microglial activation: systematic review of animal experiments. J Neuroinflammation. Jun 06 2015;12:114. doi:10.1186/s12974-015-0332-6

23. Munster BC, Aronica E, Zwinderman AH, Eikelenboom P, Cunningham C, Rooij SE. Neuroinflammation in delirium: a postmortem case-control study. Rejuvenation Res. Dec 2011;14(6):615–22. doi:10.1089/rej.2011.1185

24. Nishiguchi T, Shibata K, Yamanishi K, et al. The bispectral electroencephalography (BSEEG) method quantifies post-operative delirium-like states in young and aged male mice after head mount implantation surgery. The Journals of Gerontology: Series A. 2024;doi:10.1093/gerona/glae158

25. Liu Y, Jia S, Wang J, et al. Endocannabinoid signaling regulates post-operative delirium through glutamatergic mediodorsal thalamus-prelimbic prefrontal cortical projection. Front Aging Neurosci. 2022;14:1036428. doi:10.3389/fnagi.2022.1036428

26. Yang M, Crawley JN. Simple behavioral assessment of mouse olfaction. Curr Protoc Neurosci. Jul 2009;Chapter 8:Unit 8.24. doi:10.1002/0471142301.ns0824s48

27. Machado CF, Reis-Silva TM, Lyra CS, Felicio LF, Malnic B. Buried Food-seeking Test for the Assessment of Olfactory Detection in Mice. Bio Protoc. Jun 20 2018;8(12):e2897. doi:10.21769/BioProtoc.2897

28. Zhang J, Gao J, Guo G, et al. Anesthesia and surgery induce delirium-like behavior in susceptible mice: the role of oxidative stress. Am J Transl Res. 2018;10(8):2435–2444.

29. Lu Y, Chen L, Ye J, et al. Surgery/Anesthesia disturbs mitochondrial fission/fusion dynamics in the brain of aged mice with postoperative delirium. Aging (Albany NY). 01 12 2020;12(1):844–865. doi:10.18632/aging.102659

30. Cunningham C, Campion S, Lunnon K, et al. Systemic inflammation induces acute behavioral and cognitive changes and accelerates neurodegenerative disease. Biol Psychiatry. Feb 15 2009;65(4):304–12. doi:10.1016/j.biopsych.2008.07.024

31. Combrinck MI, Perry VH, Cunningham C. Peripheral infection evokes exaggerated sickness behaviour in pre-clinical murine prion disease. Neuroscience. 2002;112(1):7–11. doi:10.1016/s0306-4522(02)00030-1

32. Wahl D, Coogan SC, Solon-Biet SM, et al. Cognitive and behavioral evaluation of nutritional interventions in rodent models of brain aging and dementia. Clin Interv Aging. 2017;12:1419–1428. doi:10.2147/CIA.S145247

33. Chowdhury AA, Gawali NB, Munshi R, Juvekar AR. Trigonelline insulates against oxidative stress, proinflammatory cytokines and restores BDNF levels in lipopolysaccharide induced cognitive impairment in adult mice. Metab Brain Dis. Jun 2018;33(3):681–691. doi:10.1007/s11011-017-0147-5

34. Wang L, Li M, Zhu C, Qin A, Wang J, Wei X. The protective effect of Palmatine on depressive like behavior by modulating microglia polarization in LPS-induced mice. Neurochemical Research. 2022/10/01 2022;47(10):3178–3191. doi:10.1007/s11064-022-03672-3

35. Ang HP, Makpol S, Nasaruddin ML, et al. Lipopolysaccharide-Induced Delirium-like Behaviour in a Rat Model of Chronic Cerebral Hypoperfusion Is Associated with Increased Indoleamine 2,3-Dioxygenase Expression and Endotoxin Tolerance. Int J Mol Sci. Jul 31 2023;24(15)doi:10.3390/ijms241512248

36. Warner-Schmidt JL, Duman RS. Hippocampal neurogenesis: opposing effects of stress and antidepressant treatment. Hippocampus. 2006;16(3):239–49. doi:10.1002/hipo.20156

37. Altman J, Das GD. Autoradiographic and histological evidence of postnatal hippocampal neurogenesis in rats. J Comp Neurol. Jun 1965;124(3):319–35. doi:10.1002/cne.901240303

38. Shah SA, Khan M, Jo MH, Jo MG, Amin FU, Kim MO. Melatonin Stimulates the SIRT1/Nrf2 Signaling Pathway Counteracting Lipopolysaccharide (LPS)-Induced Oxidative Stress to Rescue Postnatal Rat Brain. CNS Neurosci Ther. Jan 2017;23(1):33–44. doi:10.1111/cns.12588

39. Ha SK, Moon E, Lee P, Ryu JH, Oh MS, Kim SY. Acacetin attenuates neuroinflammation via regulation the response to LPS stimuli in vitro and in vivo. Neurochem Res. Jul 2012;37(7):1560–7. doi:10.1007/s11064-012-0751-z

40. Martins S, Fernandes L. Delirium in elderly people: a review. Front Neurol. 2012;3:101. doi:10.3389/fneur.2012.00101

41. Yamanashi T, Marra PS, Crutchley KJ, et al. Mortality among patients with sepsis associated with a bispectral electroencephalography (BSEEG) score. Sci Rep. Jul 09 2021;11(1):14211. doi:10.1038/s41598-021-93588-9

42. Kozak W, Conn CA, Kluger MJ. Lipopolysaccharide induces fever and depresses locomotor activity in unrestrained mice. Am J Physiol. Jan 1994;266(1 Pt 2):R125–35. doi:10.1152/ajpregu.1994.266.1.R125

43. Godbout JP, Chen J, Abraham J, et al. Exaggerated neuroinflammation and sickness behavior in aged mice following activation of the peripheral innate immune system. FASEB J. Aug 2005;19(10):1329–31. doi:10.1096/fj.05-3776fje

44. Matarredona ER, Talaverón R, Pastor AM. Interactions Between Neural Progenitor Cells and Microglia in the Subventricular Zone: Physiological Implications in the Neurogenic Niche and After Implantation in the Injured Brain. Front Cell Neurosci. 2018;12:268. doi:10.3389/fncel.2018.00268

45. Battista D, Ferrari CC, Gage FH, Pitossi FJ. Neurogenic niche modulation by activated microglia: transforming growth factor beta increases neurogenesis in the adult dentate gyrus. Eur J Neurosci. Jan 2006;23(1):83–93. doi:10.1111/j.1460-9568.2005.04539.x

46. Tang Y, Le W. Differential Roles of M1 and M2 Microglia in Neurodegenerative Diseases. Mol Neurobiol. Mar 2016;53(2):1181–1194. doi:10.1007/s12035-014-9070-5

47. Hart BL. Biological basis of the behavior of sick animals. Neurosci Biobehav Rev. Summer 1988;12(2):123–37. doi:10.1016/s0149-7634(88)80004-6

48. Martin SA, Pence BD, Greene RM, et al. Effects of voluntary wheel running on LPS-induced sickness behavior in aged mice. Brain Behav Immun. Mar 2013;29:113–123. doi:10.1016/j.bbi.2012.12.014

49. Radulescu CI, Cerar V, Haslehurst P, Kopanitsa M, Barnes SJ. The aging mouse brain: cognition, connectivity and calcium. Cell Calcium. Mar 2021;94:102358. doi:10.1016/j.ceca.2021.102358

50. Yeh CF, Huang WH, Lan MY, Hung W. Lipopolysaccharide-Initiated Rhinosinusitis Causes Neuroinflammation and Olfactory Dysfunction in Mice. Am J Rhinol Allergy. May 2023;37(3):298–306. doi:10.1177/19458924221140965

