## Supplemental Material for "LPS-induced delirium-like behavior and microglial activation in mice correlate with bispectral electroencephalography (BSEEG)"

**eFigure 1: ……………………………………………..Page 5**

**eFigure 2: ……………………………………………..Page 6**

**eFigure 3: ……………………………………………..Page 7**

**eFigure 4: ……………………………………………..Page 8**

**eFigure 5: ……………………………………………..Page 9**

**eMethods**

The skull was exposed under isoflurane (1-3% inhalation) anesthesia. The head mount (Pinnacle Technology, Inc., Lawrence, KS; Cat# 8201) was attached with its center at the midline of the skull, its anterior holes at 2 mm anterior to bregma, and its posterior holes at 2 mm anterior to lambda ± 2 mm. Four holes were bored into the skull with a 23-gauge needle through the holes of the head mount. Four screws (Pinnacle Technology, Inc., Lawrence, KS; anterior: Cat # 8209, posterior: Cat # 8212) were attached to the skull. Finally, the head mount was fixed with dental cement. Mice received postoperative analgesia with meloxicam (VET one, Boise, ID; NDC# 13985-559-10) 5mg/kg via subcutaneous injection. After a 2-week recovery period, mice were connected by wire to an EEG system (Pinnacle Technology, Inc., Lawrence, KS; Cat# 8200-K1-SL). EEGs were recorded from the right frontal cortex using the right anterior screw attached to the skull (designated EEG2) and from the parietal cortex using the right posterior screw attached to the skull (designated EEG1). A left frontal cortex lead using the left anterior screw was attached to the skull as a ground, and a left parietal cortex lead using the left posterior screw was attached to the skull as a reference. We analyzed signals from the electrode EEG2 lead for BSEEG scoring based on the results of our previous study, which showed EEG2 to be more sensitive than EEG1^1,2^.

**Additional reference**

1.	Yamanashi T, Malicoat JR, Steffen KT, et al. Bispectral EEG (BSEEG) quantifying neuro-inflammation in mice induced by systemic inflammation: A potential mouse model of delirium. *J Psychiatr Res*. Jan 2021;133:205-211. doi:10.1016/j.jpsychires.2020.12.036

2.	Nishiguchi T, Shibata K, Yamanishi K, et al. The bispectral electroencephalography (BSEEG) method quantifies post-operative delirium-like states in young and aged male mice after head mount implantation surgery. *The Journals of Gerontology: Series A*. 2024;doi:10.1093/gerona/glae158

**eTable 1. All reagents or resources**

| Antibodies | | |
| --- | --- | --- |
| CD68 (E3O7V) Rabbit mAb | Cell Signaling Technology | Cat# 97778 |
| Goat anti-Rabbit IgG (H+L) Highly Cross-Adsorbed Secondary Antibody, Alexa Fluor™ Plus 594 | Invitrogen | Cat# A32740 |
| Anti Iba1, Goat | FUJIFILM Wako Pure Chemical Corporation | Cat# 011-27991 |
| Donkey anti-Goat IgG (H+L) Cross-Adsorbed Secondary Antibody, Alexa Fluor™ 488 | Invitrogen | Cat# A11055 |
| Chemicals | | |
| 0.9% Sodium Chloride Injection, USP | Hospira | NDC# 0409-4888-02 |
| Lipopolysaccharides from Escherichia coli O111:B4 | Sigma-Aldrich | Cat# L2630 |
| Meloxicam | VET one | NDC# 13985-559-10 |
| PBS (10X), pH 7.4 | Gibco-Thermo Fisher Scientific | Cat# 70011044 |
| Paraformaldehyde 16% Aqueous Solution EM Grade | Electron Microscopy Sciences | Cat# 15710 |
| Bovine Serum Albumin, lyophilized powder, ≥98% (agarose gel electrophoresis) | Sigma-Aldrich | Cat# A3733 |
| VECTASHIELD Vibrance® Antifade Mounting Medium with DAPI | Vector Laboratories | Cat# H-1800 |
| Software and algorithms | | |
| SMART 3.0 BASIC PACK | Panlab Harvard Apparatus | Cat# 76-0695 |
| Sirenia® software | Pinnacle Technology | N/A |
| Web-based BSEEG score calculator | Our laboratory original | N/A |
| Other | | |
| 2 EEG/1 EMG Mouse Headmount | Pinnacle Technology | Cat# 8201 |
| 0.10” EEG Mouse Screws | Pinnacle Technology | Cat# 8209 |
| 0.12” EEG Mouse Screws | Pinnacle Technology | Cat# 8212 |
| 3-Channel EEG/EMG Mouse System | Pinnacle Technology | Cat# 8200-K1-SL |
| Open Field Apparatus | Stoelting | Cat# 10-000-294 |
| Y Maze Variant F/Mice | Panlab Harvard Apparatus | Cat# 76-0079 |
| HM 450 Fully Automated Sliding Microtome | Epredia | N/A |
| All-in-one fluorescence microscope BZ-X800 | KEYENCE | N/A |

**eTable 2. Z scores on all mice**

| Saline | BFT:  Latency to pellet | OFT:  Latency to center | OFT:  Freezing time | OFT:  Time spent in center | Y maze:  Entries in novel arm | Y maze:  Duration in novel arm | Composite Z score |
| --- | --- | --- | --- | --- | --- | --- | --- |
| No 1 | -1.9 | 0.42 | 0.88 | 1.33 | -1.4 | -1.01 | -0.83 |
| No 2 | -0.28 | 0.98 | 0.47 | 1.07 | -1.4 | -0.75 | 0.04 |
| No 3 | 1.29 | 0.28 | 0.35 | -1.71 | 0.07 | -1.37 | -0.54 |
| No 4 | -0.8 | -2.22 | 1.06 | -0.03 | -0.67 | -0.28 | -1.45 |
| No 5 | 1.45 | 0.63 | -1.08 | 0.87 | 0.21 | -0.25 | 0.9 |
| No 6 | 1.14 | -0.27 | 0.17 | 0.07 | 0.8 | -0.08 | 0.91 |
| No 7 | -0.12 | 0.63 | -0.22 | 0.73 | 0.36 | 1.93 | 1.63 |
| No 8 | -0.22 | 1.18 | -2.48 | -0.69 | 1.68 | 1.2 | 0.33 |
| No 9 | 0.33 | 0.16 | 0.15 | 0.41 | -0.37 | -0.48 | 0.1 |
| No 10 | -0.5 | -0.98 | 0.14 | -1.3 | -0.52 | 0.08 | -1.52 |
| No 11 | -0.41 | -0.79 | 0.55 | -0.74 | 1.24 | 1 | 0.42 |
|  |  |  |  |  |  | Mean | 0 |
| LPS 0.5 mg/kg | BFT:  Latency to pellet | OFT:  Latency to center | OFT:  Freezing time | OFT:  Time spent in center | Y maze:  Entries in novel arm | Y maze:  Duration in novel arm | Composite Z score |
| No 1 | 3.29 | 0.21 | -1.59 | 0.41 | 0.51 | -2.03 | 0.39 |
| No 2 | 2.25 | -0.29 | -0.47 | 2.13 | 0.21 | -0.83 | 1.48 |
| No 3 | -0.74 | 1.48 | -0.42 | -1.64 | 0.36 | 0.03 | -0.46 |
| No 4 | 2.61 | 1.9 | -1.8 | 4.06 | 1.24 | 0.08 | 4 |
| No 5 | 0.79 | 1.71 | -1.04 | -0.94 | -0.37 | -1.77 | -0.8 |
| No 6 | 1.58 | 2.18 | -2.54 | 1.98 | 1.39 | 0.36 | 2.45 |
| No 7 | 1.35 | 10.99 | -3.58 | 2.31 | -0.52 | -1.99 | 4.24 |
| No 8 | 4.14 | 0.75 | -1.87 | -1.6 | 1.1 | 0.51 | 1.5 |
| No 9 | 2.01 | 4.76 | -2.14 | 1.27 | 1.54 | -0.15 | 3.6 |
| No 10 | 0.29 | 0.26 | -1.26 | -1.03 | 0.8 | 0.69 | -0.12 |
| No 11 | 1.85 | 1.61 | -4.05 | 1.6 | 2.27 | -2.18 | 0.54 |
|  |  |  |  |  |  | Mean | 1.53 |
| LPS 1.0 mg/kg | BFT:  Latency to pellet | OFT:  Latency to center | OFT:  Freezing time | OFT:  Time spent in center | Y maze:  Entries in novel arm | Y maze:  Duration in novel arm | Composite Z score |
| No 1 | 2.5 | 4.21 | -2.73 | 5.6 | 2.57 | 2.72 | 7.35 |
| No 2 | 0 | 1.9 | -2.06 | -0.57 | 0.65 | 0.41 | 0.16 |
| No 3 | 1.88 | 0.28 | -2.01 | 0.68 | 1.68 | 0.34 | 1.41 |
| No 4 | 2.5 | 10.83 | -3.99 | 1.6 | 2.57 | 1.83 | 7.58 |
| No 5 | 0.45 | 0.67 | -2.89 | 2.46 | 2.42 | 1.12 | 2.09 |
| No 6 | 1.96 | 6.08 | -1.54 | 0.77 | 0.51 | 0.13 | 3.91 |
| No 7 | 1.38 | 1.35 | -0.28 | 1.22 | 1.68 | 0.63 | 2.96 |
| No 8 | 2.87 | 7.64 | -2.94 | -0.61 | 2.42 | 2.54 | 5.89 |
| No 9 | 1.95 | 11.06 | -3.81 | 1.91 | 3.59 | 1.2 | 7.86 |
| No 10 | 2.81 | 5.56 | -2.47 | 3.93 | 1.24 | -2.18 | 4.4 |
| No 11 | 1.41 | 0.49 | -0.01 | 0.15 | -0.08 | -0.46 | 0.74 |
|  |  |  |  |  |  | Mean | 4.03 |

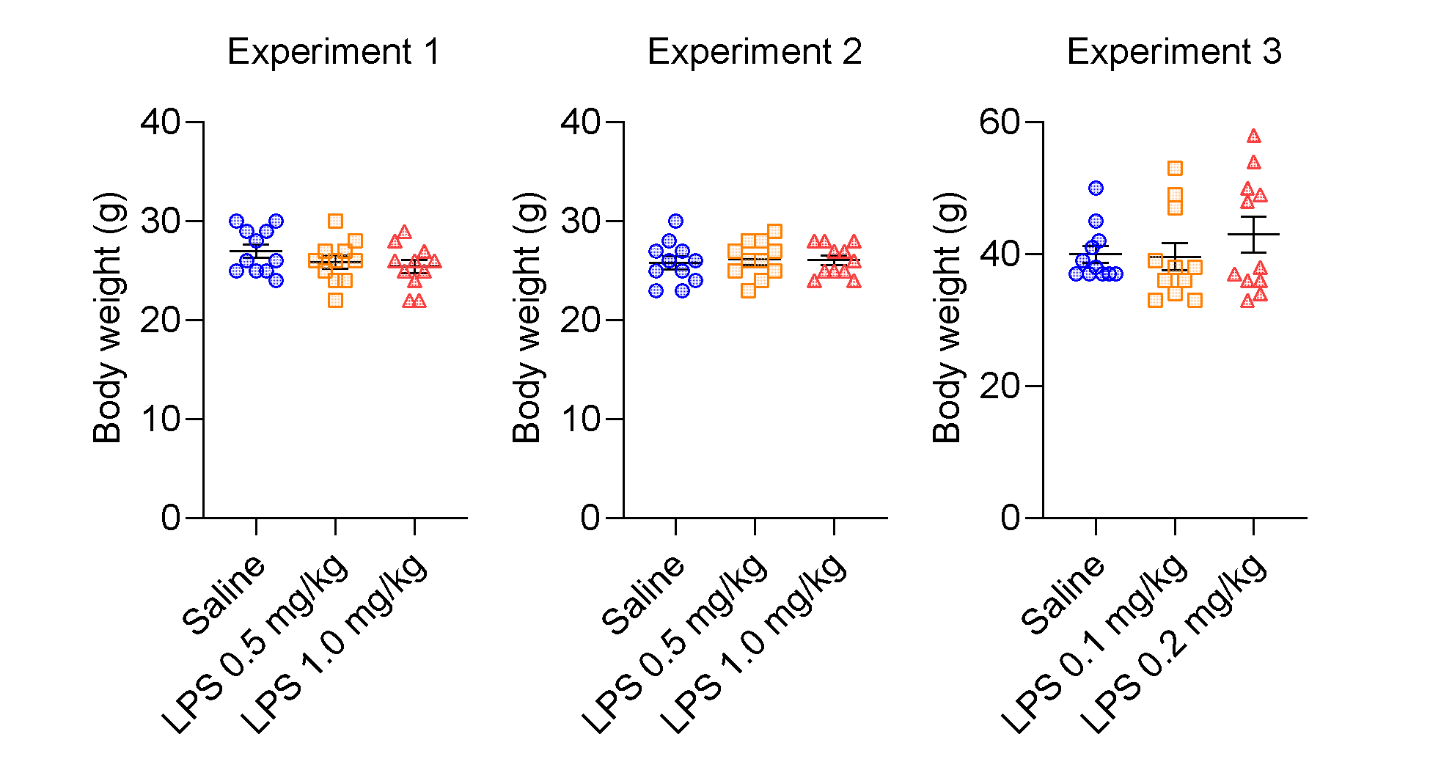

**eFigure 1. Body weight after the head mount implantation surgery.**

Experiment 1; Saline, n=11; LPS 0.5 mg/kg, n=11; LPS 1.0 mg/kg, n=11; Saline vs. LPS 0.5 mg/kg: p = 0.76, Saline vs. LPS 1.0 mg/kg: p = 0.33, LPS 0.5 mg/kg vs. LPS 1.0 mg/kg: p = 1.00; mean: Saline = 27, LPS 0.5 mg/kg = 25.91, LPS 1.0 mg/kg = 25.45. Experiment 2; Saline, n=11; LPS 0.5 mg/kg, n=11; LPS 1.0 mg/kg, n=11; Saline vs. LPS 0.5 mg/kg: p = 1.00, Saline vs. LPS 1.0 mg/kg: p = 1.00, LPS 0.5 mg/kg vs. LPS 1.0 mg/kg: p = 1.00; mean: Saline = 25.82, LPS 0.5 mg/kg = 26.18, LPS 1.0 mg/kg = 26.09. Experiment 3; Saline, n=11; LPS 0.1 mg/kg, n=11; LPS 0.2 mg/kg, n=11; Saline vs. LPS 0.1 mg/kg: p = 1.00, Saline vs. LPS 0.2 mg/kg: p = 1.00, LPS 0.1 mg/kg vs. LPS 0.2 mg/kg: p = 1.00; mean: Saline = 40, LPS 0.1 mg/kg = 39.64, LPS 0.2 mg/kg = 43.00.

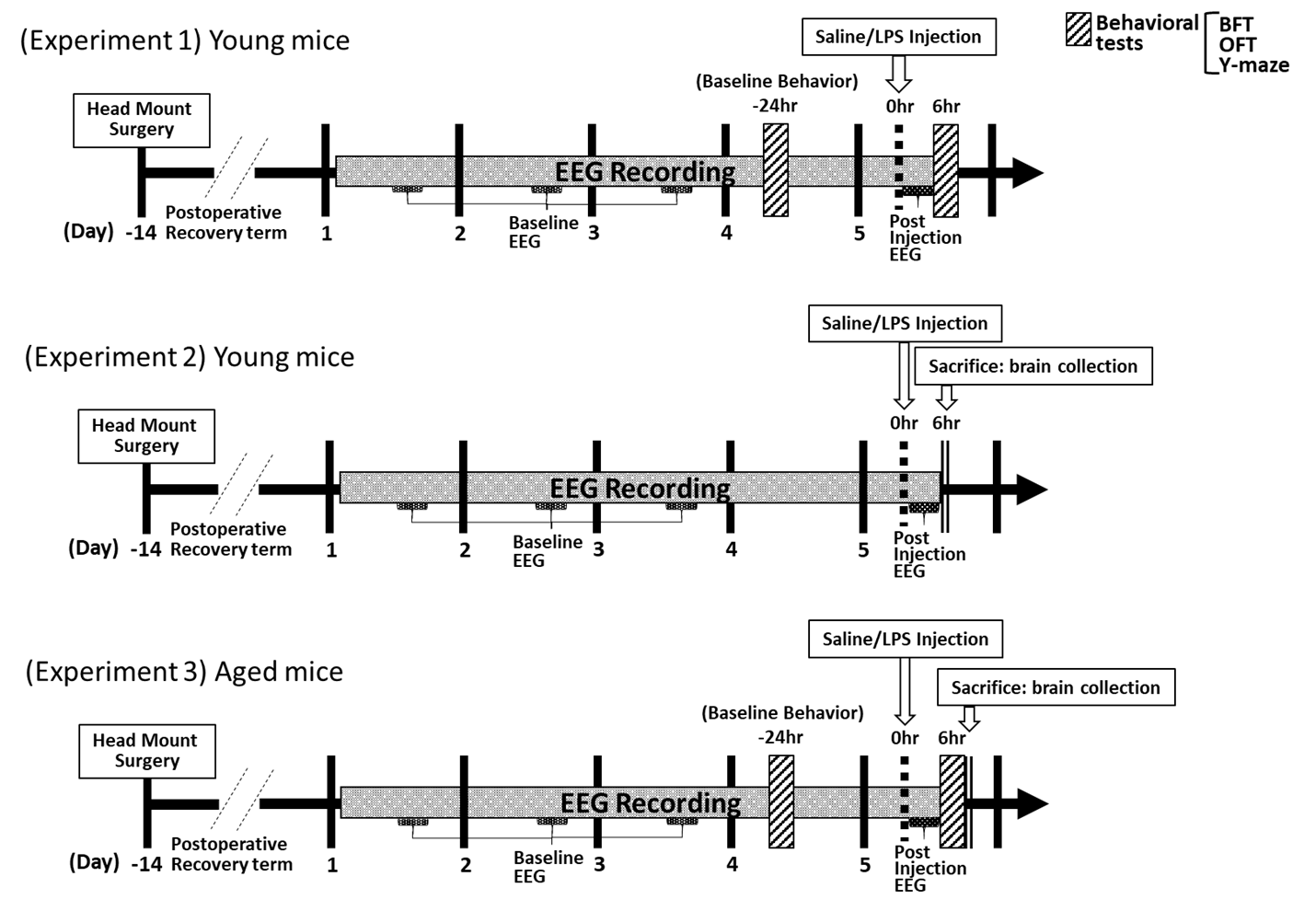

**eFigure 2. Experiment Schedules.**

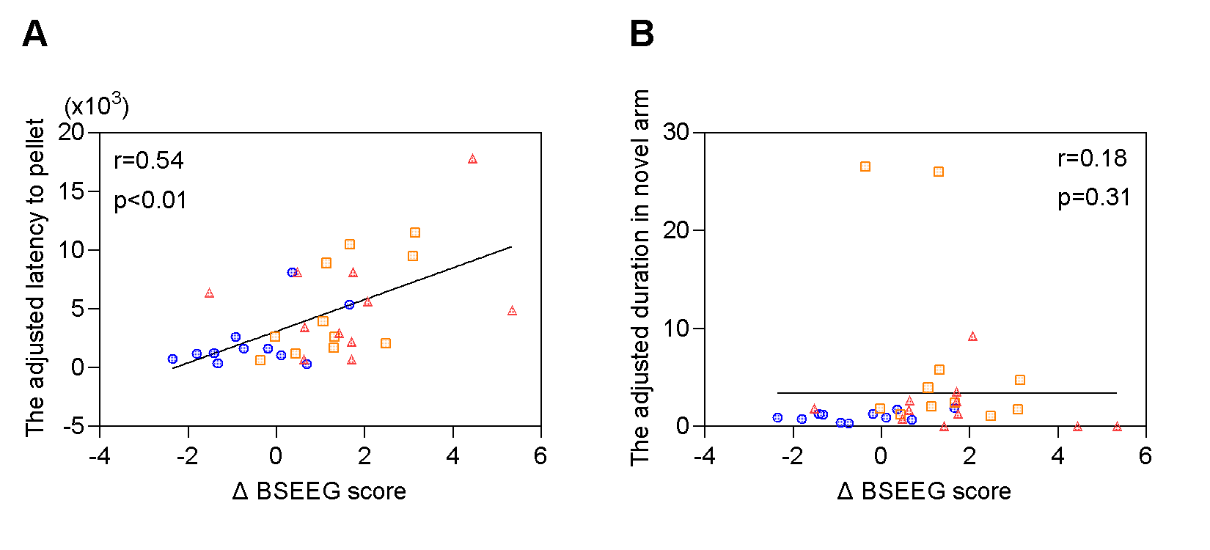

**eFigure 3. The correlation between each parameter adjusted by the total distance in OFT with Δ BSEEG score (Experiment 1)** **(Saline, n=11; LPS 0.5 mg/kg, n=11; LPS 1.0 mg/kg, n=11).**

(A) The correlation between latency to pellet in BFT adjusted by total distance in OFT and Δ BSEEG score.

(B) The correlation between duration in novel arm in Y maze test adjusted by total distance in OFT and Δ BSEEG score.

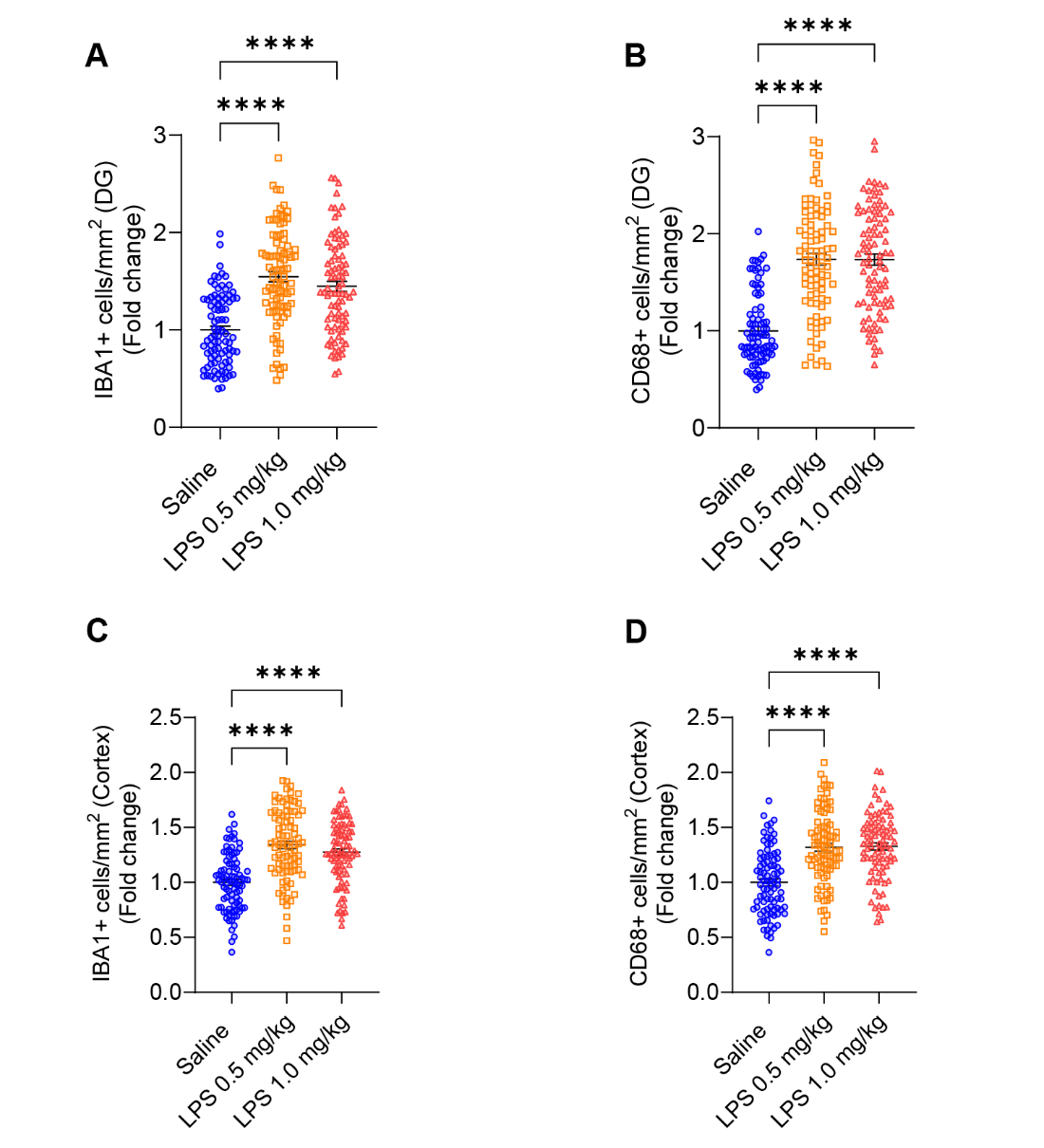

**eFigure 4. Brain slice images (****88/group) cell counts in hippocampal DG and cortex 6 hours after LPS i.p. injection in young mice (Experiment 2) (Saline, n=11; LPS 0.5 mg/kg, n=11; LPS 1.0 mg/kg, n=11) (2 (right and left) images/slice x 4 slices/mouse).**

(A) IBA1 positive cell counts in images of brain slices from hippocampal DG (Saline vs. LPS 0.5 mg/kg: p < 0.0001, Saline vs. LPS 1.0 mg/kg: p < 0.0001).

(B) CD68 positive cell counts in images of brain slices from hippocampal DG (Saline vs. LPS 0.5 mg/kg: p < 0.0001, Saline vs. LPS 1.0 mg/kg: p < 0.0001).

(C) IBA1 positive cell counts in images of brain slices from cortex (Saline vs. LPS 0.5 mg/kg: p < 0.0001, Saline vs. LPS 1.0 mg/kg: p < 0.0001).

(D) CD68 positive cell counts in images of brain slices from cortex (Saline vs. LPS 0.5 mg/kg: p < 0.0001, Saline vs. LPS 1.0mg/kg: p < 0.0001).

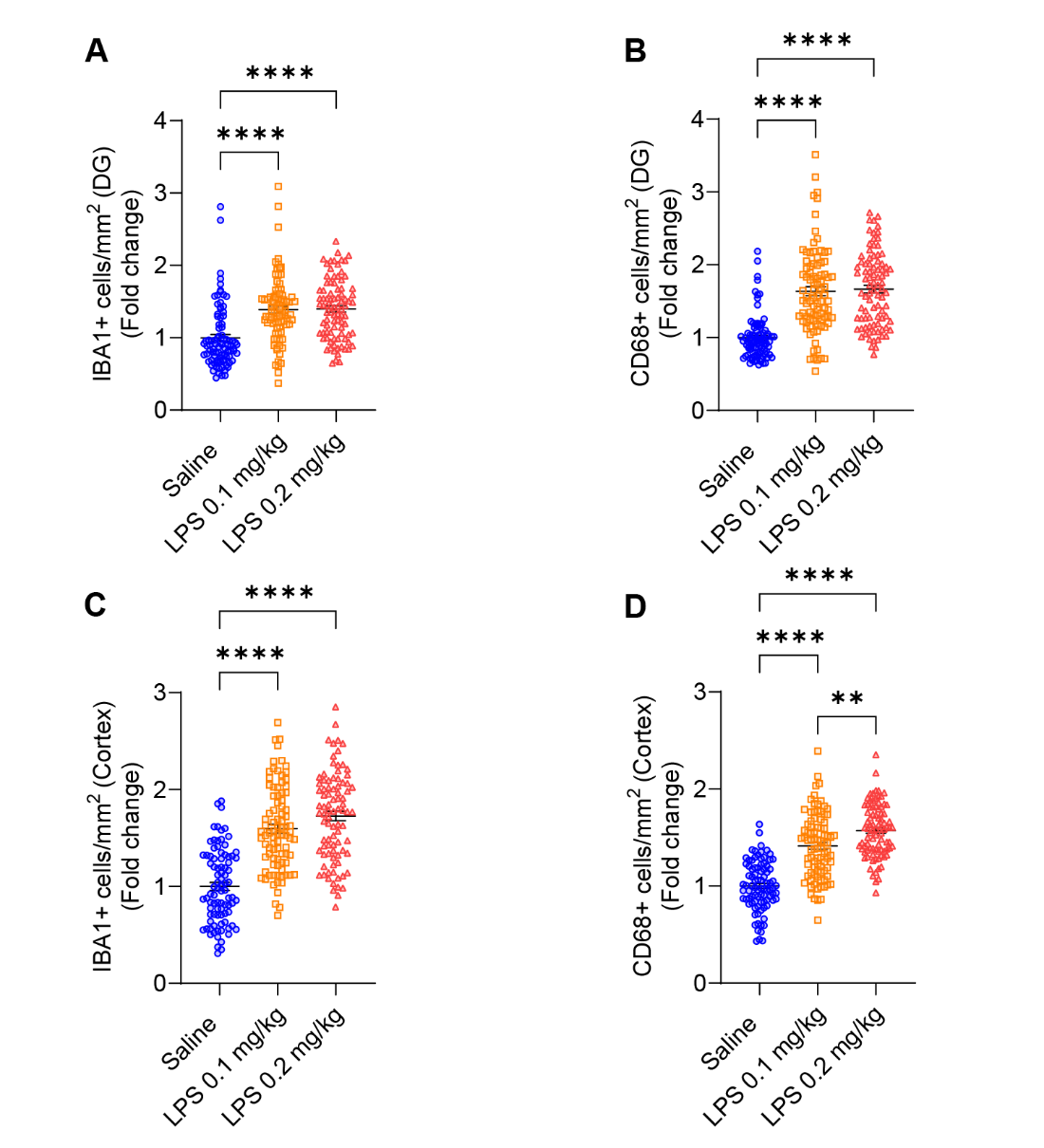

**eFigure 5. Brain slice images (88/group) cell counts in the hippocampal DG and cortex 6 hours after LPS i.p. injection in aged mice (Experiment 3) (Saline, n=11; LPS 0.1 mg/kg, n=11; LPS 0.2 mg/kg, n=11) (2 (right and left) images/slice) (4 slices/mouse).**

(A) IBA1 positive cell counts on brain slice images in the hippocampal DG (Saline vs. LPS 0.1 mg/kg: p < 0.0001, Saline vs. LPS 0.2 mg/kg: p < 0.0001).

(B) CD68 positive cell counts on brain slice images in the hippocampal DG (Saline vs. LPS 0.1 mg/kg: p < 0.0001, Saline vs. LPS 0.2 mg/kg: p < 0.0001).

(C) IBA1 positive cell counts on brain slice images in the cortex (Saline vs. LPS 0.1 mg/kg: p < 0.0001, Saline vs. LPS 0.2 mg/kg: p < 0.0001).

(D) CD68 positive cell count on brain slice images in the cortex (Saline vs. LPS 0.1 mg/kg: p < 0.0001, Saline vs. LPS 0.2 mg/kg: p < 0.0001, LPS 0.1 mg/kg vs. LPS 0.2 mg/kg: p = 0.0064).
